# The mechanism of artemisinin resistance of *Plasmodium falciparum* malaria parasites originates in their initial transcriptional response

**DOI:** 10.1101/2021.05.17.444396

**Authors:** Lei Zhu, Rob W. van der Pluijm, Michal Kucharski, Sourav Nayak, Jaishree Tripathi, François Nosten, Abul Faiz, Chanaki Amaratunga, Dysoley Lek, Elizabeth A Ashley, Frank Smithuis, Aung Pyae Phyo, Khin Lin, Mallika Imwong, Mayfong Mayxay, Mehul Dhorda, Nguyen Hoang Chau, Nhien Nguyen Thanh Thuy, Paul N Newton, Podjanee Jittamala, Rupam Tripura, Sasithon Pukrittayakamee, Thomas J Peto, Olivo Miotto, Lorenz von Seidlein, Tran Tinh Hien, Hagai Ginsburg, Nicholas PJ Day, Nicholas J. White, Arjen M Dondorp, Zbynek Bozdech

## Abstract

The emergence and spread of artemisinin resistant *Plasmodium falciparum*, first in the Greater Mekong Subregion (GMS), and now in East Africa, is a major threat to global malaria eliminations ambitions. To investigate the artemisinin resistance mechanism, transcriptome analysis was conducted of 577 *P. falciparum* isolates collected in the GMS between 2016-2018. A specific artemisinin resistance-associated transcriptional profile was identified that involves a broad but discrete set of biological functions related to proteotoxic stress, host cytoplasm remodeling and REDOX metabolism. The artemisinin resistance-associated transcriptional profile evolved from initial transcriptional responses of susceptible parasites to artemisinin. The genetic basis for this adapted response is likely to be complex.

**One sentence summary:** The transcriptional profile that characterize artemisinin resistant infections with malaria parasites *Plasmodium falciparum* originates in the initial transcriptional response to the drug.

## Introduction

Artemisinin based combination therapies (ACTs) have been critical to the success in reducing the global burden of falciparum malaria between 2000 and 2015(*1*). Loss of these drugs to resistance would be a disaster. Historically the Greater Mekong Subregion (GMS) has been the origin of antimalarial drug resistance in *P. falciparum*. In recent years the emergence and spread of artemisinin resistance and, subsequently, partner drug resistance has led to high failure rates of ACTs in several parts of the GMS(*2, 3*). Recently artemisinin resistance has arisen independently in East Africa (parts of Rwanda and Uganda)(*4*). The phenotypic manifestation of artemisinin resistant *P. falciparum* infections *in vivo* is slowed parasite clearance after treatment with artesunate. The slow clearance phenotype, defined by a parasite clearance half-life (PC½) >5hr, is attributed to loss of sensitivity of *P. falciparum* to artemisinins during the early stage of the intraerythrocytic developmental cycle (IDC), ring stage(*5, 6*) and is causally associated with nonsynonymous mutations in the propeller region of the *P. falciparum kelch 13* gene (*PfK13*)(*7, 8*). Artemisinin resistance was first reported in western Cambodia, Pailin province in 2009. Initially over 20 different *PfK13* mutations were associated with the slow parasite clearance phenotype(*8-10*). However, since 2013 the soft sweeps resulting in multiple emergences of *PfK13* mutations in the eastern part of the GMS were largely replaced by a hard selective sweep of a haplotype bearing a single nonsynonymous *PfK13* SNP (C580Y)(*11*). This single lineage of artemisinin resistant *P. falciparum* spread and expanded through western and northern Cambodia, northeastern Thailand and southern Vietnam and Lao PDR(*8, 12, 13*). This was soon joined with molecular markers associated with resistance to the ACT partner drug piperaquine. PfK13-propeller domain mutations in artemisinin resistant parasite lineages have also emerged independently and spread through Myanmar and western Thailand(*14*). Moreover, PfK13-propeller domain mutations have been reported in Northern India(*15*), and more recent foci include independent emergence in Papua New Guinea(*16*), Rwanda(*4*), Ethiopia (*17*) and other parts of sub-Saharan Africa (*18*).

The molecular mechanism by which the *PfK13* mutations confer artemisinin resistance is a subject of intense research using *in vitro* and *in vivo* models (reviewed in(*19-23*)). Collectively, these studies have proposed involvement of multiple cellular and metabolic processes in artemisinin resistance including haemoglobin degradation, proteotoxic/unfolded protein stress response, vesicular biogenesis as well as oxidative stress response and mitochondrial functions. Translational suppression mediated by phosphorylation of eIF2α, linked to “dormancy” (cell quiescence) and slowing of the IDC can also confer artemisinin resistance in *in vitro*(*24*). Missense or loss-of-function alleles of other genes were also shown to contribute to artemisinin resistance *in vitro*. These include coronin(*25*), falcipain2a/b(*26*), ubiquitin hydrolase (*pcubp1*)(*27*), and μ-subunit of the AP2 vesicular trafficking complex (*pcap2*)(*27*). Thus, it appears that artemisinin resistance is mediated by a multifaceted mechanism that results from a concerted action of several metabolic and cellular factors. These may be in different, not mutually exclusive, combinations(*28*). Undoubtedly these mechanisms drive artemisinin resistance of *P. falciparum* in *in vitro* conditions in which the parasites are supplied with superfluous amounts of nutrients, kept at uniform temperature, are not targeted by the host’s immune system and other ambient stresses exerted by the host’s environment(*29*). The question now is what are the roles of each of these identified components of artemisinin resistance in natural infections, *in vivo*.

To investigate this, several genome-wide association studies (GWAS) were conducted. These identified large regions on chromosome 10 and 13 and 14(*30, 31*), and subsequently seven nonsynonymous SNPs associated with *PfK13* SNPs(*32*). A concurrent longitudinal study of the GMS parasites collected between 2001-2014 suggested that additional genes might be associated with artemisinin resistance including an additional *kelch* protein on chromosome 10(*33*). Collectively, these studies detected genes that could be loosely linked with the artemisinin resistance-implicated biological functions, however, no experimental evidence of their role in the artemisinin resistance clinical phenotype has been reported so far.

Transcriptome-wide association analysis (TWAS) is currently emerging as the method of choice for identifying causative genetic variations of complex traits in a wide range of biological systems ranging from plants(*34*) to human(*35*). This is based on the wealth of GWAS studies showing that the vast majority of genetic polymorphisms associated with complex genetic traits (such as genetic diseases) lay within the noncoding regions(*36*). These are typically affecting DNA regulatory sequences and thus gene expression through which the phenotype is manifested(*37*). This likely also applies to *P. falciparum* as suggested by our earlier study investigating expression quantitative trait loci (eQTL) in the TRACI parasite isolates(*38*). We then carried out a TWAS of the *P. falciparum* parasites from the Tracking Resistance to Artemisinin Collaboration (TRACI) study conducted in the GMS countries between 2011-2013(*8, 39*). This showed that artemisinin resistance is associated with broad transcriptional changes of many genes, some of which may be linked with inductions of the unfolded protein response (UPR) and with a general deceleration of the IDC. Here we present a TWAS of artemisinin resistance using parasite isolates from a more recent cohort of *P. falciparum* natural infections collected in the GMS between 2016-2018; 5-7 years after the initial TRACI study(*8*), and after the recent selective sweep of *PfK13* C580Y(*2*). This uncovered a spectrum of transcriptionally correlated genes that likely contribute to artemisinin resistance via their altered transcriptional levels. We termed this the artemisinin resistance-associated transcriptional profile (ARTP) and provide evidence that its constitutive expression may have evolved from the initial transcriptional responses of sensitive *P. falciparum* parasites to the artemisinins.

## Results

### Transcriptome of the *P. falciparum* population in the GMS 2016-2018

The main purpose of this study was to identify specific genes, presumably acting in concordance with the *PfK13* mutations, whose expression activity mediates/contributes to the physiological state that enables the parasite cell to withstand the parasiticidal effects of artemisinins. We conducted TWAS of *P. falciparum* parasites derived from the blood of patients with uncomplicated *P. falciparum* infections in 13 field sites across 6 GMS countries (**Fig. 1a and Table S1**). These samples were collected during a large clinical treatment trial (TRACII) carried out from 2016 to 2018 that also characterized the spread of artemisinin resistance in the GMS(*2*). We isolated total *P. falciparum* RNA from the patients’ blood samples and performed both DNA microarray and Next Generation Sequencing (RNA-seq) analysis, as previously described(*40*). Overall, we analyzed 577 samples collected at study enrollment to characterize the baseline transcriptomic profiles (baseline, ^*(bl)*^*0hr* sample set) (**Fig. S1a**). For 459 (of the 577) patients, samples were also collected 6 hours after administering an ACT to characterize the transcriptional response of *P. falciparum* parasites to the artemisinin *in vivo* (transcriptional response, ^*(tr)*^*6hr* sample set) (**Fig. S1b)**. While the DNA microarray was used to generate the transcriptomes for all collected samples, there were sufficient levels of parasite mRNA to perform RNA-seq-based transcriptome analyses for 188 and 159 of the ^*(bl)*^*0hr* and ^*(tr)*^*6hr* samples respectively(*40*) (**Fig. S1c and S1d)**.

**Figure1.**
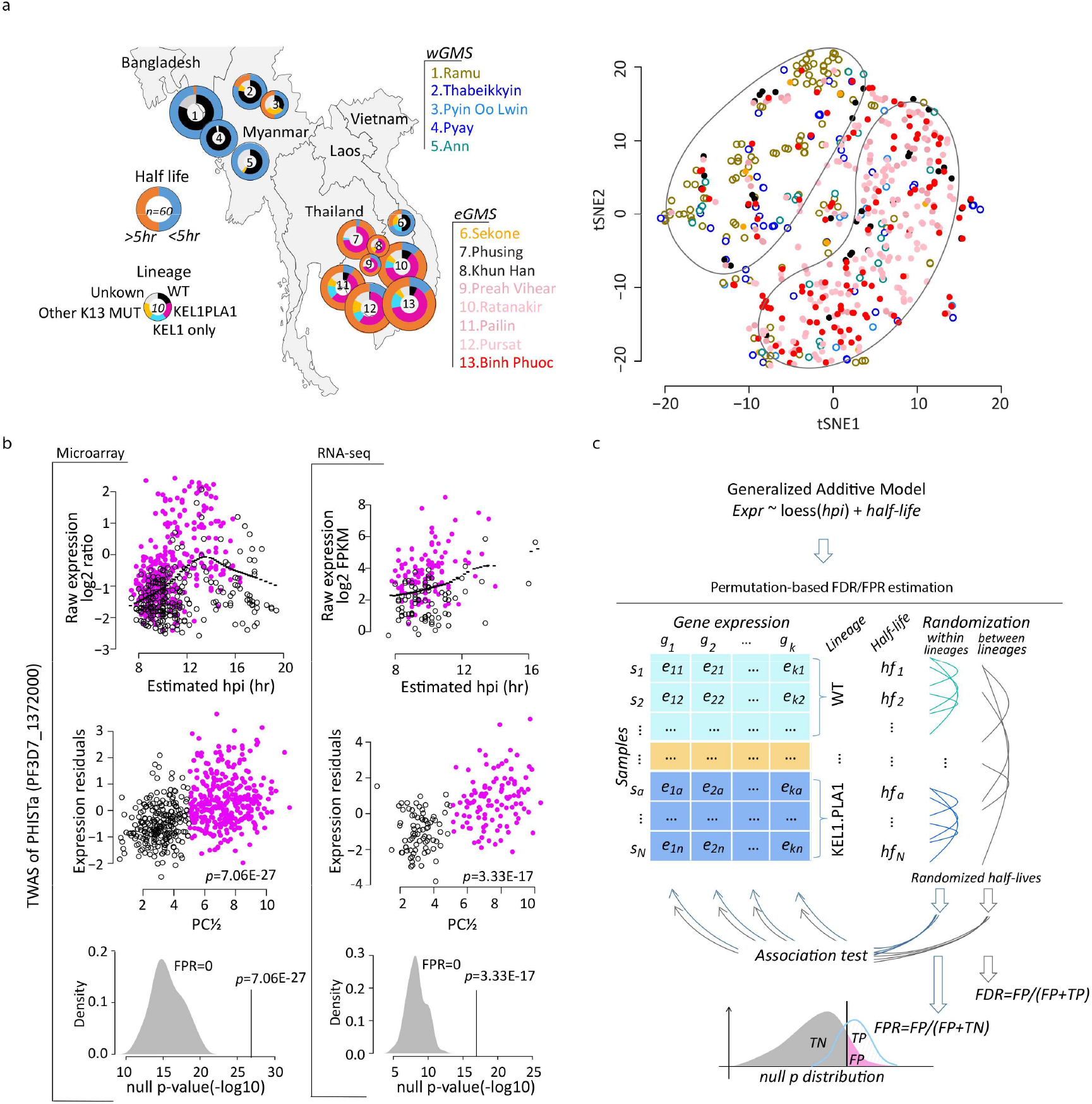
Transcriptome of the TRACII *P*.*falciparum* parasite population and a schematic illustration of the TWAS methodology. **a**. Geographic distribution of all samples used in this study. Pie charts represent the proportion of slow (PC½>5hr) and fast (PC½<5hr) clearing parasites, or lineages in categories based on *PfK13* mutations and plasmepsin II/III copy numbers (WT, KEL1 only, KEL1PLA1, other *PfK13* MUT and unknown). On the right, 577 samples before drug treatment are plotted to display two main clusters formed by the eGMS or wGMS parasites geographical distribution using t-SNE algorithm based on the top 2-12PCs. **b**. TWAS explanation using an example gene of PHISTa (PF3D7_1372000). The expression-hpi/age relationship is shown as the raw expression level (log2 ratio) plotted against the estimated hpi for the microarray data on the left (577 samples) and RNA-seq data on the right (188 samples) with purple dots representing resistant parasites (PC½>5hr), black circles for susceptible parasites (PC½<5hr) and black dotted lines for the loess curve using all dots. The expression-resistance relationship is represented by the expression residuals plotted against the PC½ for each data set. The density plots at the bottom represent 100 times permutation result within lineages for the FPR calculation. **c**. Workflow of FDR and FPR estimation for the TWAS. The null *p* distribution was built using permutated resistance status (PC½ values) across parasite samples within lineages for FPR estimation and between lineages for multiple testing correction.

From the ^*(bl)*^*0hr* sample set, we established the distribution of the *Plasmodium* life cycle developmental stages in the peripheral blood including the asexual IDC stages expressed as hours post invasion (hpi) and the fractions of sexual stages (gametocytes) as described previously(*38*) (**Fig. S2 and Data S1**). The ^*(bl)*^*0hr* sample set represented well-synchronized ring stage parasites between 6-20 hpi, with ≤ 5% of the overall parasitemia corresponding to gametocytes. PCA of the ^*(bl)*^*0hr* transcriptome revealed parasite hpi composed the major transcriptome difference across samples as it strongly correlated (Spearman *rho* = 0.87) with the first principal component (PC1) accounting for up to 32% of the overall transcriptome variation (**Fig. S3a** *top left panel*). A significant non-linear relationship was observed between the hpi and the expression (p<0.001) of most of the studied genes measured by DNA microarrays (70%) and RNA-seq (62%, **Fig. S4**). Contrast to the PC1, none of the subsequent top 2 to 12 PCs (each contributing >1% of the overall transcriptome variation) correlated with any epidemiological variables (patient sex, age and time of collection etc.) and molecular biology parameters (RNA isolation yield and quality easements etc.) suggesting that the generated dataset has minor (if any) “batch effects” from the methodological approach (**Fig. S3a** *right panel*). The t-distributed stochastic neighbor embedding (t-SNE) method using the PC2-12 revealed two major parasite groups which separated according to the geographic regions: western GMS (wGMS) and eastern GMS (eGMS) (**Fig. 1a**). This grouping pattern also corresponds to the prevalence of artemisinin resistance in the eGMS demarcated by the expansion of the *P. falciparum* lineage carrying the *PfK13* C580Y mutantion (named KEL1 lineage or PfPailin) of which 76% also carried multiple copies of plasmepsin II (KEL1PLA1, **Fig. 1a**)(*2*). *PfK13* wild type parasites from the eGMS did not exhibit any strong association with either group, which further strengthen this model (**Fig. S3a** *lower left panel*). Altogether these results suggest that the selective sweep of the “PfPailin” or “KEL1PLA1” lineage(s) in the eGMS over the last 5-7 years(*41*) is mirrored by transcriptional convergence with the eGMS parasites that is distinct from the wGMS. This transcriptome pattern is presumably either contributing to the resistance mechanism and/or alleviating a fitness cost to support its selection.

### Transcriptome-wide association study of artemisinin resistance

Next, we conducted TWAS analysis on the ^*(bl)*^*0hr* samples in order to identify genes whose steady state mRNA levels correlated with the level of artemisinin resistance represented by PC½. For this we applied a generalized additive model to relate each gene’s expression to the PC½ with a loess function along hpi (example in **Fig. 1b**, upper panels). In this model, the expression residuals reflect the relative mRNA abundance unaffected by hpi and thus can be directly correlated to PC½(*2*) (**Fig. 1b** *middle panels*). To control for multiple testing in correlating between mRNA levels and PC½ values for the whole transcriptome, we calculated the false discovery rate (FDR) for each gene based on 477,000 permutations (**Fig. 1c**). The expression-resistance association could also represent an expression-lineage relationship since the artemisinin resistance status is confounded with K13 lineages that 77% of the susceptible (PC½<5hr) samples were from the WT parasites in wGMS and 95% of the resistant (PC½<5hr) samples were from the K13 mutant in eGMS (**Fig. 1a**). Due to this homogeneity, expression-resistance associations will have a lineage effect if we use the entire data set for TWAS analysis or lose power if we test only within sub geographical region (w/eGMS). To overcome this, we estimated false positive rates (FPR) for each gene to control the type I error by repeating the analysis 100 times with randomly generated permutations (**Fig. 1b** *low panels*), a method commonly used in human genetics studies(*42, 43*). In each permutation, the lineage structure was maintained and PC½ values were randomized amongst the parasites within each lineage. Subsequently FPR was calculated based on the null p distribution reflecting the probability of expression-resistance association caused by expression-lineage relationship. And FPR<0.05 (95% confidence) was applied to define robust expression-resistance associations beyond parasite lineage effect. By this we aimed to eliminate expression-lineage-associations and identified transcripts whose expression levels are associated strictly with the PC½. One of the clearest examples was PHISTa (PF3D7_1372000) which displayed a strong association with artemisinin resistance at p=7.06E-27 (FDR=0) with FPR=0 (**Fig. 1b**, for more examples see **Fig. S5**).

Next, we applied the above TWAS method to the DNA microarray-derived data of 577 ^*(bl)*^*0hr* samples and RNA-seq-derived data of 188 ^*(bl)*^*0hr* samples separately. We observed good correlation (Spearman *rho*=0.68) between the TWAS results obtained from these two platforms (**Fig. S6**). Merging the two results, we identified 69 upregulated and 87 downregulated transcripts whose levels were significantly associated with artemisinin resistance (FDR<0.05, corresponding to *p*<1e-10 in microarray, *p*<1e-6 in RNA-seq, *note: these criteria were used to subsequently used to define the ARTP*, **Fig. 2a**). Out of these, 60S ribosomal protein gene, L35ae (PF3D7_1142600), PHISTa (PF3D7_1372000) erythrocyte membrane protein PfEMP3 (PF3D7_0219000), and PF3D7_1328400, BEM46 (PF3D7_0818600) and PF3D7_1012000 showed the strongest correlations with PC½ for up- and downregulated genes, respectively. Overall, both the upregulated and downregulated genes contribute to a broad but defined set of biological functions previously linked with artemisinin resistance such as protein and REDOX metabolism, digestive vacuole-and mitochondrial-linked biological functions, host cell remodeling etc. (**Data S2**, also see below). When compared to the transcripts identified by the same approach in parasites collected during TRACI (between 2011-2013)(*39*), we observed a significant overlap (binomial test *p*<1e-9) in upregulated and downregulated genes, including the PHISTa, KAHRP, FIKK, ATG5 and FKBP35 (for full list see **Data S2**). For further analysis we term the transcriptional variations of the 156 selected genes (69 upregulated and 87 downregulated) the <u>a</u>rtemisinin resistance-associated transcriptional <u>p</u>rofile (ARTP) and investigate further its biological relevance for the putative artemisinin resistance mechanism.

**Figure2.**
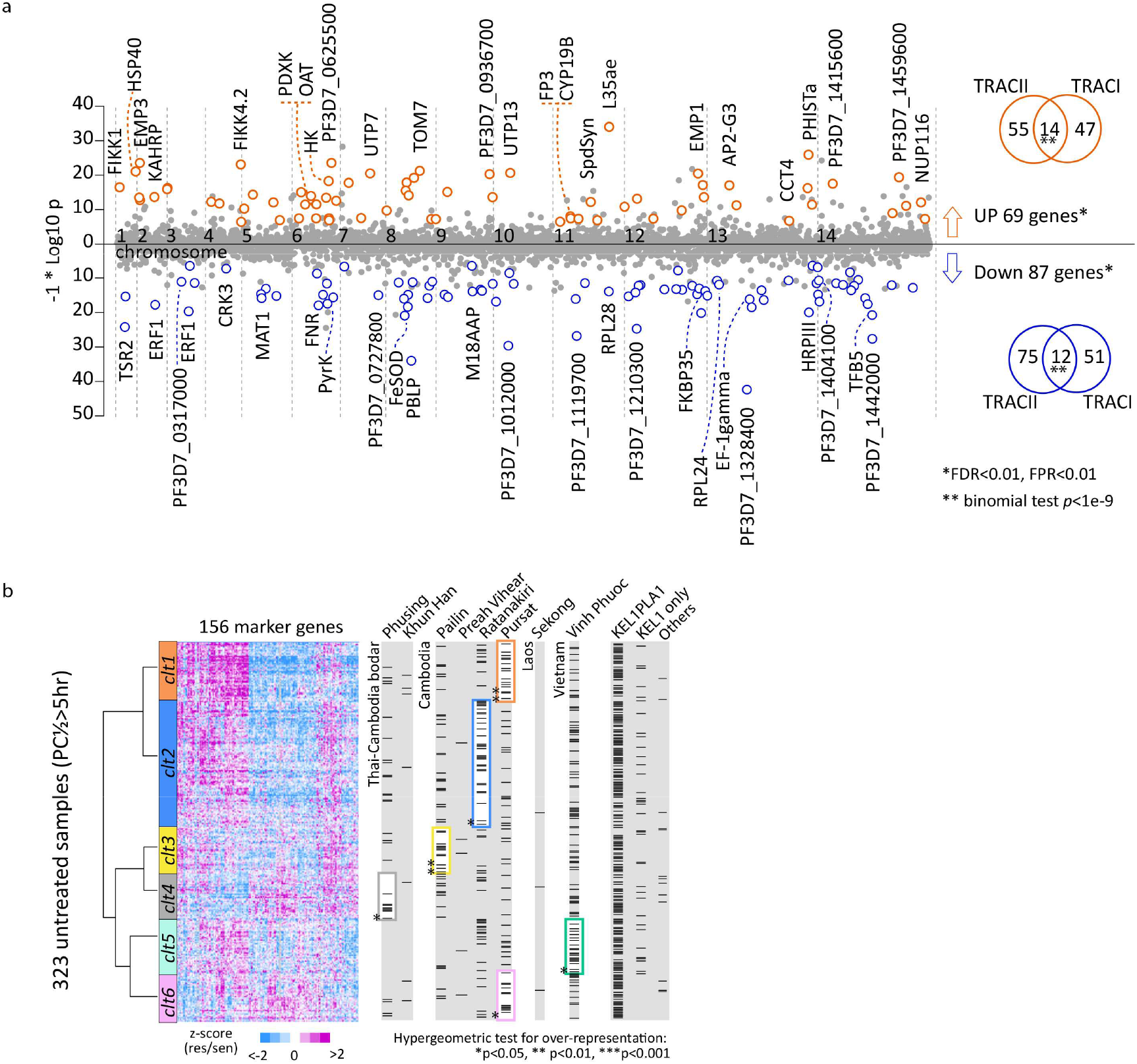
Transcriptional resistance markers. a. The scatter plot represents all the studied genes along the genomic coordinates on the X-axis with their resistance-association *p*-values displayed as -log10 *p* on the Y-axis. Genes passing the threshold (FDR<0.05, FPR<0.05) are highlighted in 69 orange circles (upregulation) and 87 blue circles (downregulation). The Venn diagrams show the overlap of TWAS results between the TRACII and TRACI studies. b. A heatmap represents the transcriptional profiles clustering for 323 eGMS ^*(bl)*^*0hr* samples showing prolonged PC½(>5hr) based on the 156 resistance markers. The colour (purple to blue) indicates the level of differential expression (upregulation to downregulation) in the resistant parasites. The left dendrogram represents Ward’s clustering result and the colour bars represent the 6 clusters obtained by clustering tree cutting. On the right, samples in each column (marked by black bars) are categorized by their respective sites or lineages. Frames mark the overrepresentations of categorized samples in the corresponding clusters. Stars indicate the statistical significance. Sample origins and index numbers are indicated for the six *PfK13* WT samples.

### Artemisinin resistance-associated transcriptional profile

Next, we aimed to investigate the utility of the ARTP as a marker of the spread and evolution of artemisinin resistance in the GMS and possibly beyond. Hence, we conducted a clustering analysis to the 323 resistant parasite samples (PC½>5hr). Specifically, the Ward’s method was applied for clustering based on the Euclidean distance matrix of similarity of the ARTP formed by the 156 transcript levels in each of the ^*(bl)*^*0hr* samples. This approach identified six clusters that displayed a distinct geographical segregation (hypergeometric test *p*<0.05) (**Fig. 2b**). It yielded two main clades with high relatedness between cluster 1 (Pursat) and 2 (Ratanakiri) on one side and the rests of the sites in cluster 3-6 including, Phusing (4) and Binh Phuoc (5) and Pailin (3). The majority of the 323 samples were from KEL1PLA1 lineage (216/323, 67%) with the remainder made up of 36 (11%) from KEL1, 19 (6%) from other lineages and 52 (17%) from unknown lineages. No lineage bias was observed with any cluster. These results suggest that the ARTP, which includes at least 156 genes with specific transcriptional “re-tuning”, could itself be a determinant of resistance to artemisinins. They also allow for the speculation that though in these resistant parasite samples, the ARTP seems to function mainly in conjunction with the nonsynonymous *PfK13* SNP, and that the parasites with this expression profile could be resistant to artemisinins even in the absence of *PfK13* mutations.

### *In vivo* transcriptional response of *P. falciparum* to ACTs

To complement the baseline TWAS, we evaluated the ^*(tr)*^*6hr* sample sets to assess transcriptional responses of *P. falciparum* exposed to ACT *in vivo*. The ^*(tr)*^*6hr* parasites exhibited a tighter distribution of the IDC stage compared to those in the ^*(bl)*^*0hr*, falling between 8-14 hpi with a median ∼12 hpi, which might reflect the more rapid loss of mature ring stage parasites because of their higher sensitivity to artemisinins (**Fig. 3a and S2**) (*authors note*). There was also a reduction (2-5%) in the fraction of gametocytes. Comparative transcriptome analysis between the ^*(bl)*^*0hr* and ^*(tr)*^*6hr* was conducted to identify transcriptional responses for the resistant (KEL/PLA1 with PC½>5hr) and susceptible (WT with PC½<5hr) parasites separately (for details see **Methods**). In total, we identified 20 and 73 genes that were induced or repressed respectively after 6hr *in vivo* treatment in the KEL1PLA1/resistant parasites. Similarly, 33 and 106 genes were induced or repressed respectively in the WT/susceptible parasites (FDR<0.05, corresponding *p*<1e-14) (**Fig. 3b**). Hence, the susceptible and resistant parasites exhibited distinct transcriptional responses to artemisinin with a minimal (albeit statistically significant) overlap including 12 and 34 commonly induced and repressed genes, respectively (**Data S2)**. Out of these, six genes were also downregulated in the baseline ^*(bl)*^*0hr* sample set while the only upregulated gene in both sample sets was PHISTa (PF3D7_1372000). There was a skewed distribution of the ^*(tr)*^*6hr* induced and repressed genes in the WT/susceptible parasites along the distribution of the transcriptional associations of the ^*(bl)*^*0hr* parasites (**Fig. 3c**). Specifically, the 33 drug-induced genes were ranked significantly towards upregulation in ^*(bl)*^*0hr* and the 106 drug-repressed genes towards downregulation (FDR=0). Contrastingly, the ^*(tr)*^*6hr* induced genes in the PfPailin (KEL1PLA1)/resistant parasites show no association with the ^*(bl)*^*0hr* baseline ARTP but the ^*(tr)*^*6hr* repressed genes exhibit a moderate overlap with upregulated genes in the ^*(bl)*^*0hr* baseline ARTP (**Fig. 3c**). This suggests that the transcriptional response of *P. falciparum* to 6 hours artemisinin exposure *in vivo* is related to the ARTP which could be its precursor (see discussion).

**Figure3.**
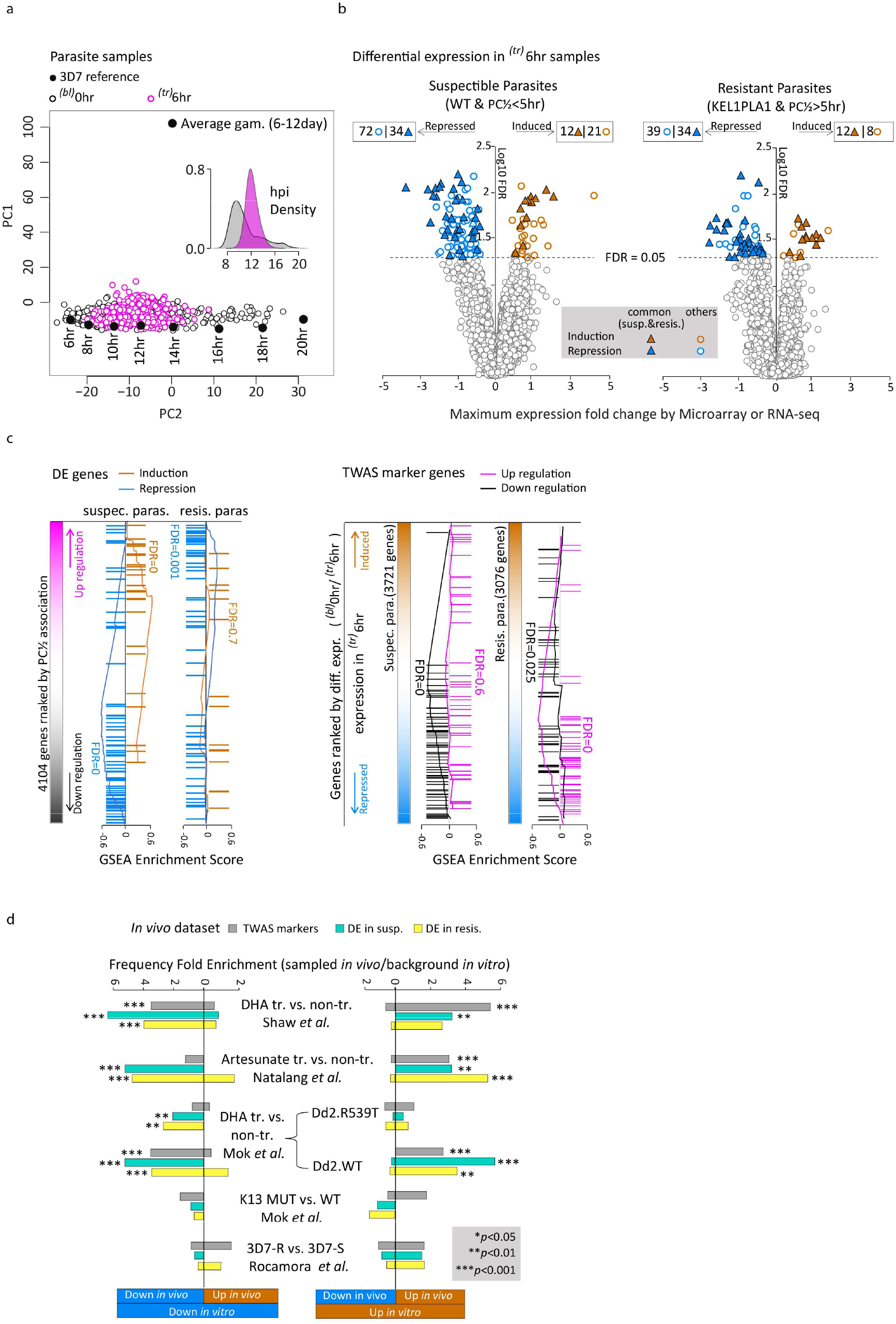
*In vivo* transcriptional response to artemisinins. **a**. The principal components space of PC1 vs. PC2 was constructed by the PCA on reference transcriptomes of the laboratory strain 3D7 at ring stage and gametocyte stages (average of the day 5^th^-12^th^). It is used to visualize transcriptome differences driven by parasite ages/hpi or developmental stages (asexual/sexual). The 577 ^*(bl)*^*0hr* (black circles) and 459 ^*(tr)*^*6hr* (purple circles) samples are projected onto this space to show their age differences. The density plot represents the estimated hpi distribution for ^*(bl)*^*0hr* (grey) and (^*tr)*^*6hr* (purple) parasites. **b**. Volcano plot represents each gene’s association *p* value of differential expression against the average expression fold change between the ^*(tr)*^*6hr* and ^*(bl)*^*0hr* parasites for the susceptible group (left, WT with PC½<5hr) and the resistant group (right, KELPLA1 with PC½>5hr) respectively. **c**. Genes differentially expressed as upregulation/induction (orange) and downregulation/repression(blue) are marked along the rank of their association to PC½ (from TWAS). Markers associated with PC½ positively (purple) or negatively(black) are marked along the rank of their differential expression levels (from b). GSEA was applied to estimate the FDR for ranking bias to either side of upregulation/induction or downregulation/repression. **d**. Bar plots represent significant overlaps between our *in vivo* study and other independent *in vitro* studies, *In vivo*: baseline TWAS analysis (^*(bl)*^*0hr*, 156 resistant markers, grey); post-treatment differentially expressed genes (^*(tr)*^*6hr*/^*(bl)*^*0hr*) in susceptible (turquoise) and resistant parasites (yellow) group, *In vitro*: transcriptional response to DHA treatment in the K1 rings(*44*); Artesunate treatment in the FCR3 strain(*45*); DHA treatment in the Dd2 R539T or WT stain(*46*); differential expression at the baseline level between the *PfK13* MUT and WT stains(*46*) as well as that between lab-derived ART-resistant and ART-sensitive 3D7 ring parasites(*47*). Stars mark intersections having >3 genes with hypergeometric test *p*<0.05.

There were highly significant gene-by-gene overlaps between the *in vivo* results and several *in vitro* studies of transcriptional responses of *P. falciparum* to artemisinin(s) (**Fig. 3d and Table 1**). This applies to the ^*(bl)*^*0hr* TWAS as well as the *in vivo* transcriptional response of ^*(tr)*^*6hr* parasites. There are good agreements between these results and earlier studies of *in vitro* transcriptional response of *P. falciparum* K1 strain exposed to dihydroartemisinin (DHA) for 1 hour(*44*), the FCR3 strain exposed to artesunate for 3 hours(*45*) and the recently reported transcriptional responses of the Dd2^wt^ strain exposed to DHA for 6 hours(*46*). This is consistent with our observation showing that the basal level ARTP overlaps well with transcriptional responses of the susceptible parasites (**Fig. 3c**). We also observed a limited overlap with our previously derived *in vitro P. falciparum* parasites with ring stage specific artemisinin resistance(*47*) suggesting that additional mechanisms conferring artemisinin resistance exist and could rise *in vivo* in the future.

**Table 1.**
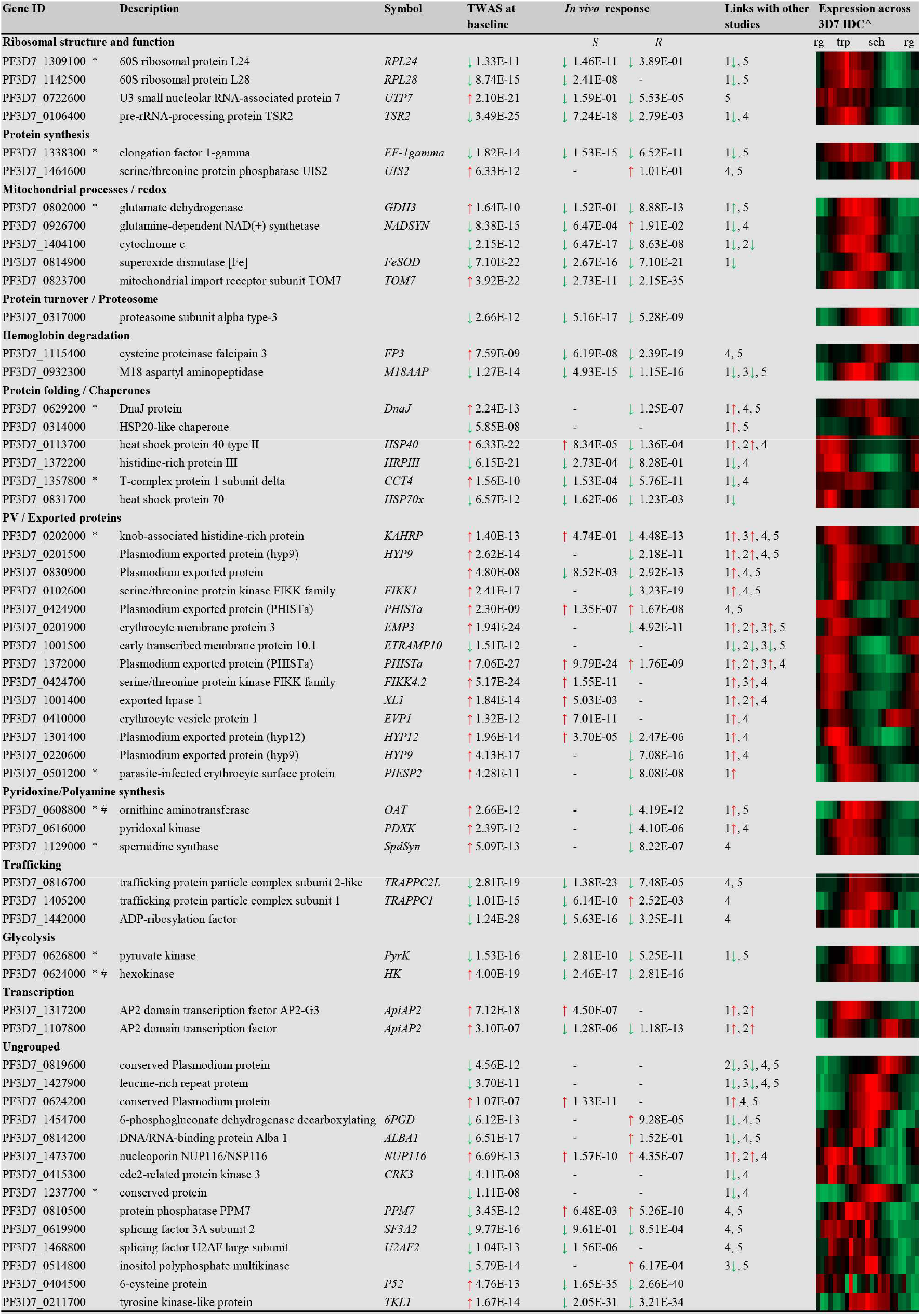
Top selected transcriptional resistance markers derived from TWAS at baseline level (^*(bl)*^*0hr*) and from *in vivo* treatment (^*(tr)*^*6hr*) sample sets. Markers are grouped according to their literature-based functional assignments and cross-referenced to other independent transcriptomics studies. Putative direct artemisinin targets are indicated(*/#). For better visualization of co-expressed transcripts relative expression levels across 3D7 IDC *in vitro* are shown in the last column (red – upregulation, green – downregulation). *Artemisinin targets by Wang, J. *et al*. (2015). #Artemisinin targets by Ismail, HM *et al*. (2016). 1) 1h DHA-treated vs. non-treated K1 at ring stage, Shaw, JP. *et al*. (2015) 2) 3h Artesunate-treated vs. non-treated FCR3, Natalang, O. *et al*. (2008) 3) Differentially expressed genes from TWAS analysis of clinical samples during TRAC1 study, Mok, S. *et al*. (2015) 4) Differentially expressed genes between K13 mutant and wild type strains Mok, S. *et al*. (2021) 5) Differentially expressed genes in in vitro Art-resistance selected 3D7 strains, Rocamora, J. *et al*. (2018). ^RNA-Seq IDC relative expression values from Kucharski, M. *et al*. (2020). R-resistant, S-sensitive, ↑ upregulation, ↓ downregulation, rg-rings, trp-trophozoites, sch-schizonts. Uncorrected p-values are indicated.

## Discussion

This study identified at least 156 genes whose altered transcription may contribute significantly to artemisinin resistance. Artemisinin resistance involves a complex array of processes that have been selected to counter the parasiticidal effect(*20-23*). These processes occur early in the asexual cycle and attenuate ring stage parasite killing. We found marked overlaps between the ARTP and the genes involved in the transcriptional response to the direct action of artemisinin observed *in vivo* (**Fig. 3a and 3b**) and/or *in vitro*(*44-47*). This suggests that the mechanisms contributing to artemisinin resistance have arisen (at least in part) from the initial transcriptional response of *P. falciparum* parasites to the direct effect of the drug in which the artemisinin induced transcriptional changes became constitutive. Indeed, there are many overlaps on the gene-by-gene level between gene expression changes in the baseline ARTP and genes induced/suppressed by artemisinins *in vivo* and *in vitro* (**Table 1**). In particular, we identified many determinants of protein metabolism including translation, folding and degradation that were also found to be a part of *P. falciparum* response to artesunate(*45*), and DHA of both susceptible and resistant parasite lines *in vitro*(*44, 46, 47*). Notable examples include gene encoding 60S ribosomal subunits (L24), and elongation factors EF-1-gamma whose proteins products were found to be direct artemisinin targets in the parasite cells(*48, 49*). Related to this, we observed significant transcriptional changes for several determinants of protein turnover and protein folding from which DnaJ proteins and the T-complex-protein 1 subunit are likely to be direct protein targets of the artemisinin drugs (**Table 1**). We also found transcriptional suppression of factors involved in REDOX functionalities, some of which are related to mitochondrial functions recently shown to play a key role in artemisinin resistance *in vitro*(*46*). There was marked transcriptional activity of genes involved in biosynthetic pathways including pyridoxine/polyamine and purine/pyrimidine synthesis and glycolysis. Interestingly several enzymes encoded by these transcripts were also found to be inhibited by artemisinin directly including ornithine aminotransferase (OAT), spermidine synthase, pyruvate kinase and hexokinase(*48, 49*). This also applies to the *P. falciparum* pyridoxal kinase, PDXK, whose mammalian ortholog, was shown to interact directly and being inhibited by artesunate(*50*). Finally, the ARTP contains strong baseline-level upregulations of several transcripts encoding proteins exported to the host erythrocyte. These are implicated in host cell remodeling and/or host-parasite interactions that are paralleled by drug-induced transcriptional responses (**Table 1**) (see below). Taken together, these observations support a model in which the initially adaptive transcriptional response became constitutively expressed as a result of drug selection, predisposing the parasite to withstand the drug’s parasiticidal effect.

Functional assignments of the “transcriptional markers” of both the ARTP and *in vivo* induced artemisinin responses revealed by this study support their role in key biological processes aligned with both *PfK13*-dependent and independent mechanism(s) of artemisinin resistance suggested by previous *in vitro* studies (summarized in **Fig. 4**). First, *PfK13*, the key causal factor in artemisinin resistance, was shown to localize predominantly to the cytostomes, possibly regulating endocytosis of hemoglobin(*51*). *PfK13* mutations lead to a reduced rate of endocytosis and thus hemoglobin digestion, which in turn lessens the bioactivation of artemisinin as a result of lower levels of Fe^2+^ in the parasite cytoplasm. We observed baseline upregulation of factors involved in hemoglobin degradation and their suppression in sensitive parasites after 6hr *in vivo* treatment. Second, reductions of the *PfK13* protein levels, resulting from the mutations, were also shown to suppress the proteotoxic shock and subsequent cell death normally induced by artemisinins(*52, 53*). This suppression can be reversed by proteasome inhibitors in both *P. falciparum*(*54*) and *P. berghei*(*55*). We found marked transcriptional suppression of protein synthesis, folding and turnover including the core subunits of the proteasome (**Data S2 and Fig. 4**). These were accompanied by transcriptional variability of factors involved in transcription, mRNA processing, ribosomal biogenesis (RiBi), translation, translational translocation and protein transport (**Table 1 and Data S2**). This is also consistent with our earlier TWAS study of the TRACI samples where we observed upregulation of the UPR t at is canonically related to proteotoxic shock response(*38, 39*). Third, PfK13 was also shown to function as an adaptor for the cullin3-RING-E3 ubiquitin ligase targeting a specific set of proteins for proteasome-mediated degradation(*56*). Mutated forms of PfK13 fail to recognize their targets including phosphoinositide 3-kinase (PI3K), which subsequently leads to an increase in PI3P vesicles that alters the rate of hemoglobin endocytosis(*57*). A set of genes involved in vesicular trafficking and the underlying lipid metabolism was also found to be transcriptionally associated with artemisinin resistance in this study (**Table 1 and Fig. 4**). Fourth, *PfK13* mutations appear to mediate rewiring of the *P. falciparum* metabolic program to higher levels of survival(*58*). This involves mainly induction of mitochondrial processes including damage sensing and oxidative stress response that counteract artemisinin induced oxidative and alkylation activity(*54*). In line with this, induced oxidative stress response was also detected in two independently derived *in vitro P. falciparum* artemisinin resistant lines(*47, 59*). One of the chief functions of the parasite mitochondria lies in regeneration of ubiquinone, a necessary ECT component in pyrimidine biosynthesis pathways(*60*) which was also shown to be altered by *PfK13* mutations(*46*). Here we observed transcriptional variations of multiple REDOX and mitochondria related factors including purine/pyrimidine synthesis pathway-related markers (OPRT, HGPRT) suppressed in susceptible parasites but also several downregulated transcripts pertaining to ECT (SDHB, CytC) and TCA-metabolism (**Data S2**). Fifth, several genes encoding proteins exported to the host cell cytoplasm were found amongst the transcriptional resistance markers. These include several members of the FIKK and PHIST gene families that are mostly implicated in host-cell remodeling affecting rigidity and/or cytoadhesion of the infected RBCs(*61-64*). This could suggest that altered host-parasite interactions might contribute to artemisinin resistance; albeit indirectly by reducing parasite clearance by the spleen. Given the functional diversification of the FIKK and PHIST as well as other *P. falciparum* gene families annotated as “exported proteins” their direct role in artemisinin resistance mechanisms cannot be ruled out. Several of the exported proteins factors as well as many of the above-mentioned biological pathways have been recently linked with artemisinin resistance in a large-scale *P. falciparum piggyBac-transposon* mutant screen(*65*). Altogether these observations support the majority of the previous studies implicating a wide spectrum of biological functions discovered by *in vitro* experiments to play specific roles in artemisinin resistance *in vivo*.

**Figure4.**
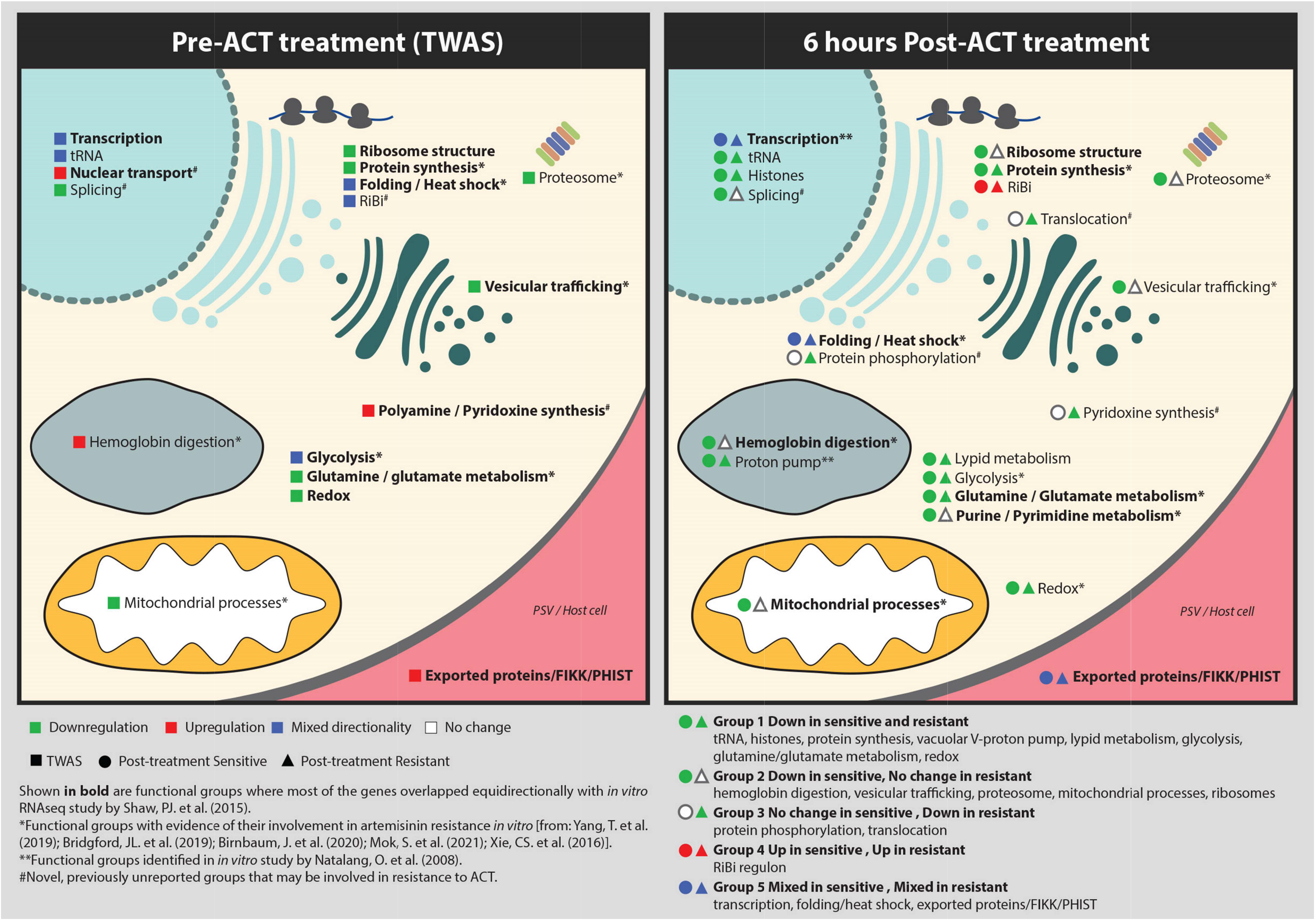
Functional assignment of resistance markers. Functional assignments of transcriptional artemisinin resistance markers derived from TWAS analysis at baseline-level (^*(bl)*^*0hr*) and that of genes repressed/induced after 6 hours of ACT treatment (^*(tr)*^*6hr*). Colors represent transcriptomics directionality for each group. The association to previous independent *in vitro* studies is also shown.

In summary, our data suggest that a characteristic transcriptional pattern (the ARTP) is emerging in the artemisinin resistant parasites in the GMS as a result of a complex adaptive response to artemisinin. The ARTP mitigates parasite damage in young asexual parasites by which it has evolved to become more efficient. In contrast to TWAS, GWAS have until now failed to identify such factors make limited observations of direct links between the molecular effectors of artemisinin resistance *in vivo;* with the exception of the *PfK13* SNP(*32*). Although it is still possible that more mutations in the coding regions may contribute to artemisinin resistance but were not detected because of typical sample size limitations of GWAS(*37*); in the context of a highly complex *P. falciparum* population structure in the GMS(*66*). Our result strongly suggests that many causative mutations might be situated within DNA regulatory elements affecting regulation of gene expression. It was previously shown that the majority of sequence polymorphisms in *P. falciparum* occurs in the noncoding regions accumulated particularly around core promoter regions as Insertion/Deletion within Short Tandem Repeats (STR)(*67*). The high AT content (∼85%) of the *P. falciparum* noncoding regions hampered most whole genome sequencing efforts on field samples done up until now. However, the results of this study make a case for synergizing GWAS and TWAS (combined into eQTL studies) as a potential strategy to monitor the spread and evolution of malaria drug resistance in the future genomic epidemiological surveys.

## Supporting information

Data S1 estimated HPI and gametocyte fraction

Data S2 selected genes list

## Acknowledgements

We would like to thank all members of the Tracking Resistance to Artemisinin Collaboration for providing access to clinical samples used in this study as well as all patients that have agreed to participate in this TRACII clinical trials. We also would like to thank Sachel Mok for providing the parasite reference pool and Cholrawee Promnarate, Nakarin Aud-Ai for coordinating field specimen collection. This research was funded by Singapore National Medical Research Council grant # NMRC/OFIRG/0040/2017 and Singapore Ministry of education grant # MOE2019-T3-1-007 and MOE2017-T2-2-030 (S) awarded to Z. Bozdech. Moreover, A. Dondorp coordinated funding for the TRACII epidemiology studies funded by DFID (FCDO): Artemisinin Resistant Malaria Research Programme (TRAC). DFID PO 5408 and Wellcome Trust: MOP Thailand Core award 220211/Z/20/Z and 220211/A/20/Z. Author contributions: Lei Zhu performed all the analysis of this study, prepared the figures and drafted the manuscript. Rob Van Der Pluijm and Zbynek Bozdech designed the study, performed the analysis and drafted the manuscript; Michal Kucharski performed all lab experiments including the design of the experimental protocols, necessary validations and helped to analyze the data and draft the manuscript and figures; Jaishree Tripathi contributed to the protocol design for lab experiments; Sourav Nayak helped to process RNA-Seq data; Dominic P. Kwiatkowski, François Nosten, Hagai Ginsburg, Nicholas PJ Day, Nicholas J. White and Arjen M. Dondorp participated in the design and supervised the project and gave vital comments to the Discussion. Tracking Resistance to Artemisinin Collaboration II authors carried out all the field and laboratory work necessary to obtain *P. falciparum* samples. The manuscript was reviewed by all the authors.

## Data and materials availability

All data is available in the main text or the supplementary materials or from the corresponding author upon request. All transcriptome data used in this study are available at NCBI’s Gene Expression Omnibus (GEO) database with the series accession number GSE149735 for microarray and GSE169520 for RNA-seq.

Needs statement on application of a CC-BY licence to ensure open access.

## References and Notes

1. T. W. H. Organization, Malaria Report 2019. (2019).

2. R. W. van der Pluijm et al., Triple artemisinin-based combination therapies versus artemisinin-based combination therapies for uncomplicated Plasmodium falciparum malaria: a multicentre, open-label, randomised clinical trial. Lancet, (2020).

3. A. P. Phyo et al., Declining Efficacy of Artemisinin Combination Therapy Against P. Falciparum Malaria on the Thai-Myanmar Border (2003-2013): The Role of Parasite Genetic Factors. Clin Infect Dis 63, 784–791 (2016).

4. A. Uwimana et al., Emergence and clonal expansion of in vitro artemisinin-resistant Plasmodium falciparum kelch13 R561H mutant parasites in Rwanda. Nat Med 26, 1602–1608 (2020).

5. K. Chotivanich et al., Laboratory detection of artemisinin-resistant Plasmodium falciparum. Antimicrob Agents Chemother 58, 3157–3161 (2014).

6. B. Witkowski et al., Reduced artemisinin susceptibility of Plasmodium falciparum ring stages in western Cambodia. Antimicrob Agents Chemother 57, 914–923 (2013).

7. F. Ariey et al., A molecular marker of artemisinin-resistant Plasmodium falciparum malaria. Nature 505, 50–55 (2014).

8. E. A. Ashley et al., Spread of artemisinin resistance in Plasmodium falciparum malaria. N Engl J Med 371, 411–423 (2014).

9. W. K. G.-P. S. Group, Association of mutations in the Plasmodium falciparum Kelch13 gene (Pf3D7_1343700) with parasite clearance rates after artemisinin-based treatments-a WWARN individual patient data meta-analysis. BMC Med 17, 1 (2019).

10. S. Takala-Harrison et al., Independent Emergence of Artemisinin Resistance Mutations Among Plasmodium falciparum in Southeast Asia. The Journal of infectious diseases, (2014).

11. M. Imwong et al., The spread of artemisinin-resistant Plasmodium falciparum in the Greater Mekong subregion: a molecular epidemiology observational study. Lancet Infect Dis 17, 491–497 (2017).

12. A. M. Dondorp et al., Artemisinin resistance in Plasmodium falciparum malaria. N. Engl. J. Med. 361, 455–467 (2009).

13. R. W. van der Pluijm et al., Determinants of dihydroartemisinin-piperaquine treatment failure in Plasmodium falciparum malaria in Cambodia, Thailand, and Vietnam: a prospective clinical, pharmacological, and genetic study. Lancet Infect Dis 19, 952–961 (2019).

14. M. Boulle et al., Artemisinin-Resistant Plasmodium falciparum K13 Mutant Alleles, Thailand-Myanmar Border. Emerg Infect Dis 22, 1503–1505 (2016).

15. N. Mishra et al., Emerging polymorphisms in falciparum Kelch 13 gene in Northeastern region of India. Malar J 15, 583 (2016).

16. O. Miotto et al., Emergence of artemisinin-resistant Plasmodium falciparum with kelch13 C580Y mutations on the island of New Guinea. PLoS Pathog 16, e1009133 (2020).

17. A. G. Bayih et al., A Unique Plasmodium falciparum K13 Gene Mutation in Northwest Ethiopia. Am J Trop Med Hyg 94, 132–135 (2016).

18. E. Kamau et al., K13-propeller polymorphisms in Plasmodium falciparum parasites from sub-Saharan Africa. The Journal of infectious diseases 211, 1352–1355 (2015).

19. C. L. Ng, D. A. Fidock, Plasmodium falciparum In Vitro Drug Resistance Selections and Gene Editing. Methods Mol Biol 2013, 123–140 (2019).

20. M. R. Rosenthal, C. L. Ng, Plasmodium falciparum Artemisinin Resistance: The Effect of Heme, Protein Damage, and Parasite Cell Stress Response. ACS Infect Dis 6, 1599–1614 (2020).

21. A. M. Talman, J. Clain, R. Duval, R. Menard, F. Ariey, Artemisinin Bioactivity and Resistance in Malaria Parasites. Trends Parasitol 35, 953–963 (2019).

22. L. Tilley, J. Straimer, N. F. Gnadig, S. A. Ralph, D. A. Fidock, Artemisinin Action and Resistance in Plasmodium falciparum. Trends Parasitol 32, 682–696 (2016).

23. J. Wang, C. Xu, Z. R. Lun, S. R. Meshnick, Unpacking ‘Artemisinin Resistance’. Trends Pharmacol Sci 38, 506–511 (2017).

24. M. Zhang et al., Inhibiting the Plasmodium eIF2alpha Kinase PK4 Prevents Artemisinin-Induced Latency. Cell Host Microbe 22, 766–776 e764 (2017).

25. A. R. Demas et al., Mutations in Plasmodium falciparum actin-binding protein coronin confer reduced artemisinin susceptibility. Proc Natl Acad Sci U S A 115, 12799–12804 (2018).

26. N. Klonis et al., Artemisinin activity against Plasmodium falciparum requires hemoglobin uptake and digestion. Proc Natl Acad Sci U S A 108, 11405–11410 (2011).

27. R. C. Henrici, D. A. van Schalkwyk, C. J. Sutherland, Modification of pfap2mu and pfubp1 Markedly Reduces Ring-Stage Susceptibility of Plasmodium falciparum to Artemisinin In Vitro. Antimicrob Agents Chemother 64, (2019).

28. L. Paloque, A. P. Ramadani, O. Mercereau-Puijalon, J. M. Augereau, F. Benoit-Vical, Plasmodium falciparum: multifaceted resistance to artemisinins. Malar J 15, 149 (2016).

29. M. LeRoux, V. Lakshmanan, J. P. Daily, Plasmodium falciparum biology: analysis of in vitro versus in vivo growth conditions. Trends Parasitol 25, 474–481 (2009).

30. I. H. Cheeseman et al., A major genome region underlying artemisinin resistance in malaria. Science 336, 79–82 (2012).

31. S. Takala-Harrison et al., Genetic loci associated with delayed clearance of Plasmodium falciparum following artemisinin treatment in Southeast Asia. Proc. Natl. Acad. Sci. U. S. A. 110, 240–245 (2013).

32. O. Miotto et al., Genetic architecture of artemisinin-resistant Plasmodium falciparum. Nature genetics, (2015).

33. G. C. Cerqueira et al., Longitudinal genomic surveillance of Plasmodium falciparum malaria parasites reveals complex genomic architecture of emerging artemisinin resistance. Genome Biol. 18, 78 (2017).

34. Y. Ma et al., Combined transcriptome GWAS and TWAS reveal genetic elements leading to male sterility during high temperature stress in cotton. New Phytol, (2021).

35. M. Wainberg et al., Opportunities and challenges for transcriptome-wide association studies. Nature genetics 51, 592–599 (2019).

36. P. M. Visscher et al., 10 Years of GWAS Discovery: Biology, Function, and Translation. Am J Hum Genet 101, 5–22 (2017).

37. E. Cano-Gamez, G. Trynka, From GWAS to Function: Using Functional Genomics to Identify the Mechanisms Underlying Complex Diseases. Front Genet 11, 424 (2020).

38. L. Zhu et al., The origins of malaria artemisinin resistance defined by a genetic and transcriptomic background. Nat Commun 9, 5158 (2018).

39. S. Mok et al., Drug resistance. Population transcriptomics of human malaria parasites reveals the mechanism of artemisinin resistance. Science 347, 431–435 (2015).

40. M. Kucharski et al., A comprehensive RNA handling and transcriptomics guide for high-throughput processing of Plasmodium blood-stage samples. Malar J 19, 363 (2020).

41. W. L. Hamilton et al., Evolution and expansion of multidrug-resistant malaria in southeast Asia: a genomic epidemiology study. Lancet Infect. Dis. 19, 943–951 (2019).

42. M. A. Pourhoseingholi, A. R. Baghestani, M. Vahedi, How to control confounding effects by statistical analysis. Gastroenterol Hepatol Bed Bench 5, 79–83 (2012).

43. H. J. Westra et al., Systematic identification of trans eQTLs as putative drivers of known disease associations. Nat. Genet. 45, 1238–1243 (2013).

44. P. J. Shaw et al., Plasmodium parasites mount an arrest response to dihydroartemisinin, as revealed by whole transcriptome shotgun sequencing (RNA-seq) and microarray study. BMC Genomics 16, 830 (2015).

45. O. Natalang et al., Dynamic RNA profiling in Plasmodium falciparum synchronized blood stages exposed to lethal doses of artesunate. BMC Genomics 9, 388 (2008).

46. S. Mok et al., Artemisinin-resistant K13 mutations rewire Plasmodium falciparum’s intra-erythrocytic metabolic program to enhance survival. Nat Commun 12, 530 (2021).

47. F. Rocamora et al., Oxidative stress and protein damage responses mediate artemisinin resistance in malaria parasites. PLoS Pathog 14, e1006930 (2018).

48. J. Wang et al., Haem-activated promiscuous targeting of artemisinin in Plasmodium falciparum. Nat Commun 6, 10111 (2015).

49. H. M. Ismail et al., Artemisinin activity-based probes identify multiple molecular targets within the asexual stage of the malaria parasites Plasmodium falciparum 3D7. Proc. Natl. Acad. Sci. U. S. A. 113, 2080–2085 (2016).

50. V. B. Kasaragod et al., Pyridoxal kinase inhibition by artemisinins down-regulates inhibitory neurotransmission. Proc. Natl. Acad. Sci. U. S. A. 117, 33235–33245 (2020).

51. J. Birnbaum et al., A Kelch13-defined endocytosis pathway mediates artemisinin resistance in malaria parasites. Science 367, 51–59 (2020).

52. C. Dogovski et al., Targeting the cell stress response of Plasmodium falciparum to overcome artemisinin resistance. PLoS Biol 13, e1002132 (2015).

53. T. Yang et al., Decreased K13 Abundance Reduces Hemoglobin Catabolism and Proteotoxic Stress, Underpinning Artemisinin Resistance. Cell Rep 29, 2917–2928 e2915 (2019).

54. J. L. Bridgford et al., Artemisinin kills malaria parasites by damaging proteins and inhibiting the proteasome. Nat Commun 9, 3801 (2018).

55. N. V. Simwela et al., Plasmodium berghei K13 Mutations Mediate In Vivo Artemisinin Resistance That Is Reversed by Proteasome Inhibition. mBio 11, (2020).

56. A. Mbengue et al., A molecular mechanism of artemisinin resistance in Plasmodium falciparum malaria. Nature 520, 683–687 (2015).

57. S. Bhattacharjee et al., Remodeling of the malaria parasite and host human red cell by vesicle amplification that induces artemisinin resistance. Blood 131, 1234–1247 (2018).

58. Sachel, Rewiring. Nature Comm, (2021).

59. L. Cui et al., Mechanisms of in vitro resistance to dihydroartemisinin in Plasmodium falciparum. Mol Microbiol 86, 111–128 (2012).

60. H. J. Painter, J. M. Morrisey, M. W. Mather, A. B. Vaidya, Specific role of mitochondrial electron transport in blood-stage Plasmodium falciparum. Nature 446, 88–91 (2007).

61. C. K. Moreira et al., The Plasmodium PHIST and RESA-Like Protein Families of Human and Rodent Malaria Parasites. PLoS One 11, e0152510 (2016).

62. A. Oberli et al., Plasmodium falciparum Plasmodium helical interspersed subtelomeric proteins contribute to cytoadherence and anchor P. falciparum erythrocyte membrane protein 1 to the host cell cytoskeleton. Cell Microbiol 18, 1415–1428 (2016).

63. A. G. Schneider, O. Mercereau-Puijalon, A new Apicomplexa-specific protein kinase family: multiple members in Plasmodium falciparum, all with an export signature. BMC Genomics 6, 30 (2005).

64. P. Ward, L. Equinet, J. Packer, C. Doerig, Protein kinases of the human malaria parasite Plasmodium falciparum: the kinome of a divergent eukaryote. BMC Genomics 5, 79 (2004).

65. C. W. Min Zhanga, Jenna Oberstallera, Phaedra Thomasa, Thomas D., Ottob, c, Debora Casandra, Sandhya Boyapalle, Swamy R. Adapa, Shulin Xua, Katrina Button-Simonsd, Matthew Mayhob, Julian C. Rayner, Michael T. Ferdig, Rays H. Y. Jiang, John H. Adams, The endosymbiotic origins of the apicoplast link fever-survival and artemisinin-resistance in the malaria parasite. bioRxiv https://doi.org/10.1101/2020.12.10.419788doi, (2021).

66. O. Miotto et al., Multiple populations of artemisinin-resistant Plasmodium falciparum in Cambodia. Nature genetics 45, 648–655 (2013).

67. A. Miles et al., Indels, structural variation, and recombination drive genomic diversity in Plasmodium falciparum. Genome Res 26, 1288–1299 (2016).

