## Supplementary Materials and Methods for "The mechanism of artemisinin resistance of *Plasmodium falciparum* malaria parasites originates in their initial transcriptional response"

**This PDF file includes:**

Materials and Methods

Table S1: Summary of samples in this study.

Figure S1: Transcriptional profiles of the studied P.falciparum parasites.

Figure S2: Distribution of the estimated age, gametocytes fraction and parasite clearance half-life.

Figure S3: PCA of the parasite transcriptome.

Figure S4: Expression level correlates to age/hpi for most of the P. falciparum genes.

Figure S5: Examples for FPR calculation.

Figure S6: Consistent results of TWAS analysis across platforms

Figure S7: Defined intensity threshold for microarray-generated transcriptome filtering.

**Other Supplementary Materials for this manuscript include the following:**

Data S1: Parasite age and gametocyte proportion estimation

Data S2: TWAS markers and drug response genes

### Materials and Methods

#### Ethics statement

All the samples used for this study were collected with written informed consent from the patients or their legal guardians. All protocols were approved by the appropriate national ethics committees or the Oxford Tropical Research Ethics Committee.

#### *In vivo* sample collection

All samples used in this study were collected from patients involved in Tracking Resistance to Artemisinin Collaboration 2 (TRAC2), a multi-site trial that took place between Aug 7, 2015, and Feb 8, 2018. The total number of 1100 patients with uncomplicated *Plasmodium falciparum* malaria were recruited in eight countries. In the presented study 680 patients' samples from 6 countries and 13 sites across Southeast Asia Greater Mekong Region were analyzed (Ramu in Bangladesh; Thabeikkyin, Pyay, Pyin Oo Lwin and Ann in Myanmar; Phusing and Khun Han in Thailand; Pailin, Pursat, Ratanakiri, and Preah Vihear in Cambodia; Sekong in Laos; Bin Phuoc in Vietnam). All details regarding samples collection, site locations, inclusion criteria, parasitemia assessment, and given treatments were published previously(*1*). In brief, samples were collected from the venous blood of malaria-infected patients at admission to the clinic (Hour 0, 0hr) and 6 hours after their respective treatment (Hour 6, 6hr). Depending on their location, patients were randomly assigned different double or triple Artemisinin Combination Therapy (ACT) as follows: Thailand, Cambodia, Vietnam, and Myanmar - dihydroartemisinin-piperaquine or dihydroartemisinin-piperaquine plus mefloquine; Cambodia - artesunate-mefloquine or dihydroartemisinin-piperaquine plus mefloquine; Laos, Myanmar, Bangladesh - artemether-lumefantrine or artemether-lumefantrine plus amodiaquine. Following collection, blood was subsequently depleted of plasma and buffy coat. Hour 0 samples were additionally purified from white blood cells (WBC) using CF11 columns using the Worldwide Antimalarial Resistance Network (WWARN) protocols. 0.2 to 0.5 mL of packed red blood cells (pRBC) from each sample was homogenized by mixing with 10 volumes of TRIzol (Invitrogen), frozen at -80oC and then shipped to Nanyang Technological University, Singapore.

**RNA isolation**

All clinical samples mixed with TRIzol were supplemented with 5 volumes of chloroform (Merck) and processed as per manufacturer's instructions to obtain phase separation. Top (aqueous) phase was mixed 1:1 (v/v) with 100% analytical grade ethanol (Merck) and RNA was extracted using the Direct-Zol-96 extraction kit (Zymo) following the manufacturer's guide. For this study, we have adopted the RNA extraction kit to a high-throughput EVO 200 robotic platform (Tecan). RNA integrity, concentration and quality were assessed as described before(*2*).

**cDNA synthesis and microarray hybridization**

For each sample 250ng of total RNA was used for subsequent reverse transcription reaction followed by 19 rounds of PCR amplification using modified Smart-Seq2 method described previously(*3*). Oligo-dT30 (IDT Asia) was used as a primer to enrich for mRNA and avoid ribosomal rRNA amplification. The amplified product was purified using Ampure XP magnetic beads (Beckman Coulter) on Microlab Nimbus robotic platform (Hamilton) and 100ng of cDNA was used for subsequent 10 rounds of amplification to generate aminoallyl-coupled cDNA for the hybridizations as described(*4, 5*). 17ul (~5µg) of each Cy-5-labeled (GE Healthcare) cDNA of patient's sample and an equal amount of Cy-3-labeled (GE Healthcare) cDNA of the reference pool were then hybridized together on our customized microarray chip using commercially available hybridization platform (Agilent) for 20 hours at 70°C with rotation at 10rpm. Microarrays were washed and immediately scanned using Power Scanner (Tecan) at 10 µM resolution and with automated photomultiplier tubes gain adjustments to balance the signal intensities between both channels. The reference pool used for microarray was a mixture of 3D7 parasite strain RNA collected every 6 hours during 48 hours of the full intraerythrocytic developmental cycle.

**RNA-sequencing**

Purified 1ng of cDNA was used to generate sequencing libraries using Illumina Nextera XT kit as described in manufacturer's protocol. Purified cDNA libraries were analyzed on the Bioanalyzer High-Sensitivity DNA chips (Agilent), subsequently pooled (20-24 samples per lane) and sequenced on Illumina HiSeq4000 platform generating 150 bp paired end reads with 110 Gb data output generated per lane.

#### *In vitro* culture

Plasmodium falciparum 3D7 strain was obtained from BEI Resources (MRA-102) and maintained in purified human pRBC in RPMI 1640 medium (Gibco) supplemented with Albumax I (Gibco) (0.25%), hypoxanthine (Sigma) (0.1 mM), Sodium bicarbonate (Sigma) (2 g/L), and gentamicin (Gibco) (50 µg/L). Cultures were kept at 37°C with 5% $CO_2$, 3% $O_2$, and 92% $N_2$. Culture media were replenished every 24 hours. Freshly washed pRBC were added to the culture when necessary. Both parasitemia and parasite morphology were assessed by microscopic examination of blood smears stained with Giemsa (Sigma).

#### Intraerythrocytic asexual developmental cycle reference time course (IDC)

IDC reference time course data were obtained from the study published previously (11). In brief, prior to time-point experiment, parasites were double-synchronized with 5% sorbitol solution to achieve a synchrony of +/- 6 hours and cultured under constant agitation. For sampling of highly synchronous parasites during the asexual life cycle, the first time point was considered as the TP1 (Time Point 1) when >95% of early ring stage parasites (approx. 4 hpi) were present in the culture. Starting from TP1, parasites were collected every 2 hours for 25 successive time points. The total of 24 time points were used here to build an asexual reference transcriptome generated using two different platforms: microarrays and RNA-seq.

#### Raw transcriptome of *P.falciparum* parasites

The transcriptome was generated by quantifying the total RNAs for each parasite sample using two-color microarray or RNA-seq or both. Additionally, control samples derived from ring stages 3D7 lab strain were quantified together with those clinical samples in microarray method. It was made up a total of 30 samples for pre-treatment and 27 samples for post-treatment.

The raw data of microarray was acquired using GenePix Pro v6.0 software (Axon Instruments). Then the signal intensities were Loess-normalized within arrays followed by quantile-normalization between samples/arrays using Limma package of R(*6*). Missing values were assigned to weak signal probes showing median foreground intensity less than 1.5-fold of the median background intensity at either Cy5 (sample RNA) or Cy3 (reference pool RNA) channel. Each gene expression was estimated as the average of log2 ratios (Cy5/Cy3) for probes representing it.

For RNA-seq, raw reads were trimmed to remove sequencing adapters, amplification primers and low-quality bases from 3' ends. HISAT2 aligner(*7*) was used to perform alignment to the *P.falciparum* genome downloaded from geneDB database in version of March 2018. BEDTools(*8*) was applied to calculate the read counts for each annotated transcript based on only uniquely mapped reads with pairs in proper orientation. At last, the transcriptional level was estimated for each gene by calculating the Fragments per kilo base per million mapped reads (FPKM) at the gene.

**Developmental stage estimation**

For each parasite sample, we estimated the asexual age (hours post-invasion, hpi) and the proportion of gametocytes using the method described previously(*9, 10*). In brief, the expression value of a gene, $E_g$, is assumed as a sum of the expression in asexual stage of hpi *hr*, denoted as $x_g(hr)$, and the average expression of sexual stages (from the 5$^{th}$ to 12$^{th}$ day during gametocytes developing), denoted as $z_g$, mixed in certain proportions. The mixture model is presented in formula as:

$$E_g = (1-\alpha)*x_g(hr) + \alpha*z_g + \varepsilon_g \qquad (1),$$

where α is the proportion of gametocytes to the total parasite count (sum of sexual and asexual parasites) and the $\varepsilon_g$ is the associated error term. $x_g(hr)$ is estimated by the reference transcriptome generated during asexual intraerythrocytic cycle in *P. falciparum* 3D7 strain which has a total of 25 time points with two hours intervals of 48 hours, accessible via GEO accession number GSE149865. To obtain a higher resolution of the asexual reference transcriptome for stages estimation, 24 time points of the data (the 9$^{th}$ time point removed due to its big dissimilarity to others) were interpolated into 240 data points by smooth splines method. $z_g$ is estimated by the identical reference gametocyte transcriptome described previously(*9*), accessible via GEO accession number GSE121505. In practice, samples were normalized by control samples across batches by adding a scaling factor for each batch before applying the prediction model. The scaling factor was estimated for each batch by calculating the average expression difference between controls and the reference at corresponding ages. Finally, we evaluated the log-likelihood values over a grid of mixtures for varying the gametocyte proportion $\alpha$, and hpi *hr*, for each sample. The results of estimated hpi and gametocytes proportion were listed in **Data S1.**

**Transcriptome filtering**

To achieve high quality data for the following analyses, we pruned the sample set according to their transcriptome heterogeneity and discarded samples with low signal intensities. In practice, outlier samples were removed if they were distinct from the majorities with <10 cohort samples at the similarity threshold for grouping. The threshold was determined by the average similarity of controls to clinical samples using Spearman's *rho*. Second, we removed samples displaying extremely low intensities (mode of the intensity <10) of Cy5 signal on the microarray (**Fig. S7**). In addition, to reduce the prediction errors (if any) affecting on the following study, we removed 2 samples having very high PC½ as 13.1 and 19.3hr (>2 times Median Absolute Deviation to the median), and 24 samples with extremely high gametocytes prediction (> 18%, 3times MAD to the median) as most of the parasite samples exhibiting a low even zero proportion of gametocytes. We found all the discarded samples at this end mostly having a low parasitemia or high human content. Finally, with the microarray method, we identified a transcriptome of 4,779 genes presented across >75% of the 577 samples pre-treatment and 4,714 genes across 459 samples post-treatment. Among the total of 577 pre-treatment samples and 467 post-treatment samples, 438 samples were paired (collected from the identical patient). With the RNA-seq method, we asked for at least 1M uniquely mapped reads in a library to call for the transcriptome. Overall, we identified the transcriptome of 4305 genes with >75% representation for 188 samples pre-treatment and 3923 genes for 159 samples post-treatment for further analyses. The data is accessible in Gene Expression Omnibus (GEO) database via the series accession number GSE149735 for microarray and GSE169520 for RNA-seq.

**PCA and population transcriptomic analysis**

Principle Component Analysis (PCA) was applied to the *(bl)0hr* transcriptome data set. We inspected the top 12 PCs for the following association analysis because each of the rest PCs contributed to <1% of the overall transcriptome variations. The top 12 PCs were tested against all the clinical and technical factors collected during sample processing to assess the potential environmental influence on the population transcriptomic structure. For the factors represented in categorical variables like sampling sites, parasite lineages and patient's gender, ANOVA test was used for assessing the statistical significance of associations. For other factors in continues/discrete variables like parasitemia, hpi and patient enrollment time etc., linear regression was used to test the statistical relationship between each factor and each PC with Spearman's *rho* calculated for each pair of variables as well. The results are visualized in **Fig. S3a**.

The regression analysis revealed that parasite age (hpi) significantly (*p*= 7.96e-283) correlated to the top PC (PC1) with showing a high Spearman's *rho* as 0.87 in the *(bl)0hr* data set by microarray and 0.85 by RNA-seq. PCA of all samples including both *(bl)0hr and (tr)6hr* resulted in similar high correlations with displaying *rho* as 0.78 in microarray and 0.76 in RNA-seq (**Fig. S3b**). It indicates that hpi is the major factor distinguishing parasites transcriptome in field. Correlation was also observed between the PC2 and the ratio of parasite to human content; the PC6 and estimated gametocytes proportion but the Spearman's *rho* was dropped to 0.53 and 0.52, respectively. These two factors (parasite/human ratio and gametocytes fraction) can interact with other experimental or environmental conditions to drive the minor (compare to hpi) differences in parasites transcriptome. For this study, we considered only hpi as the major factor contributing to gene expression variation across nature parasites and used it as a major predictor variable in modeling gene expression for the further regression analysis. The potential for expression variation caused by other factors, like sampling site and parasite lineage, are analyzed for particular genes associated to resistance in the following studies.

To visualize the parasite population structure in a two-dimensional map, t-distributed stochastic neighbor embedding was applied to the PC2-12 with PC1 excluded to minimize the hpi effects. It was implemented in R with the M3C package(*11*).

**The PC space projection**

We performed PCA to the reference transcriptome of 3D7 parasite stain at ring stages together with the average transcriptome of 3D7 mature gametocytes ($5^{th}$ to $12^{th}$ day during development). The resulted PC1 clearly distinguishes the sexual and asexual stages as well as the PC2 reflects the ring-stage parasite development as it obviously separates the 8 ring-stage transcriptomes with 2hr difference in between. With the space of PC1 vs PC2, parasite age can be visualized without bias by transcriptome projection. Therefore, we normalized all the clinical samples to the center transcriptome derived from the above PCA and rotated it according to the PC1 and PC2 loadings. At last, the 577 pre-treatment samples and 467 post-treatment samples were projected onto the PC space showing an obvious age window shift in the parasites after drug treatment (**Fig. 3a**).

**Transcriptome-wide association study (TWAS)**

TWAS was carried out in order to call for the marker candidates whose mRNA levels were positively or negatively associated with artemisinin resistance. Since parasite age contributed the most to expression variation across clinical samples, we designed a generalized additive model with the age/hpi specified in a loess function to test the expression-resistance association over the dynamic relationship between expression and age. The regression analysis formula for each gene is represented as:

$$E = \beta_0 + \beta_1 * f(hpi) + \beta_2 * hf + \varepsilon \qquad (2),$$

Where $E$ denotes gene expression across samples, $f(hpi)$ denotes the function of $hpi$ which is loess regression here, $hf$ denotes the variable of PC½, $\beta_{0,1,2}$ represents the parameters of intercept and slopes to predict and $\varepsilon$ is the error terms. Therefore, the resulted p-value of this regression analysis reflect the relationship between expression residuals of age fitting to the half-lives (PC½), alternatively the expression-resistance association independent of age.

With this approach, we tested the expression-resistance association for all the genes, 4,779 genes with microarray and 4,714 genes with RNA-seq, individually. To correct for the multiple testing, we estimated FDR for each gene by permutation method which constructed a null p distribution by 477,000 times testing on the association of expression to randomized PC½ values (**Fig. 1c**). In addition, to address the cofounding relation between artemisinin resistance (PC½>5hr) and parasite genetic lineage which also largely coincided with the geographical region (w/eGMS), we estimated a FPR value for each gene to control the type I error by 100 times permutations. In each permutation, the lineage structure was maintained and PC½ values were randomized amongst the parasites within each lineage. The resulted null *p*-values reflect the significance of expression-lineage associations independent of PC½ and the FPR calculated from the null p distribution (**Fig. 1b**) reflect the probability of expression-resistance association caused by expression-lineage relationship. At last, FPR<0.05 (95% confidence) was applied to define the robust expression-resistance associations beyond parasite lineage effect. We plotted the expression residuals against the original and randomized PC½ values for PHISTa gene (shown in **Fig. 1b**) to illustrate the randomization procedure, also for three example genes with significant expression-resistance associations (FDR<0.05) and different levels of FPR (0.01, 0.53 and 1) together with other two genes with FDR>0.5 and FPR<0.01 for reference **(Fig. S5)**.

We applied this approach to microarray and RNA-seq data separately. The results agreed to each other with showing a high correlation (Pearson correlation coefficient=0.68) of the average expression fold change of resistant/susceptible between the two datasets (**Fig. S6**). We merged the markers defined by microarray or RNA-seq excluding 10 genes which displayed conflicting directions of expression changes in resistant parasite in two techniques. Finally, our TWAS approach determined 69 genes upregulated and 87 genes downregulated in the resistant parasites at FDR<0.05 (corresponding *p*<1e-10 in microarray, *p*<1e-6 in RNA-seq) and FPR<0.05.

We applied the same TWAS pipeline to the TRACI data and reanalyzed 824 samples collected during 2011-1013. It resulted in 61 expression upregulation and 63 downregulation associated to artemisinin resistance with the identical criteria above (FDR<0.05&FPR<0.05, corresponding *p*<8.5e-6). This TWAS result significantly overlaps that from TRACII with 14 upregulation and 12 downregulation in common (binomial test *p*<1e-9).

**ARTP clustering**

To investigate the resistance-associated transcriptome structure, we first defined the artemisinin resistance-associated transcriptional profiles (ARTP) for each parasite sample using the 156 marker genes. To obtain expression levels with hpi effects maximumly reduced, we second extracted out the expression residuals from the formula (2) with the hpi function fitting only for each gene. The expression residuals were normalized for the 323 resistant parasite samples

(PC½>5hr and from eGMS) against that of 104 susceptible samples (PC½<5hr and from wGMS) by calculating z-scores to represent the number of standard deviations by which the expression of the studied gene in resistant parasite was above or below the mean in susceptible parasites. Next, Euclidean distance was applied to the similarity matrix of ARTP to construct the sample distance matrix and the Ward's method was used to obtain the dendrogram of clustering tree for the 323 resistant parasite samples (**Fig. 2b**). The six clusters shown in **Fig. 2b** was defined by the tree cutting at the 1/6 of tree height using the "cutree" function in R stats package.

#### *In vivo* transcriptional response measurement

The transcriptional response to artemisinin was inspected for the field *P.falciparum* parasites by comparing the $^{(tr)}$*6hr* sample set to the $^{(bl)}$*0hr* sample set. Grouping the $^{(bl)}$*0hr* samples by lineage and resistance status level (PC½ greater/smaller than 5hr) revealed two largest groups which are KEL1PLA1 parasites with PC½>5hr (37% of the total 577 samples) and *PfK13* WT parasites with PC½<5hr (29% of 577). Excluding the lineage unknown samples, all the rest groups contain <6% of the total samples. For this study, we performed the comparative transcriptional analysis specifically for the *PfK13* WT samples with PC½<5hr (the susceptible parasites) and the KEL1PLA1 samples with PC½>5hr (the resistant parasites).

We adjusted the above regression model of formula (2) to better present the data of $^{(tr)}$*6hr* and $^{(bl)}$*0hr* samples as:

$$E = \beta_0 + \beta_1 * f(hpi) + \beta_2 * treatment + \beta_3 * resistance\_status + \varepsilon \qquad (3),$$

Where *treatment* indicates patient treatment condition which is pre-treatment ($^{(bl)}$*0hr*) or post-treatment ($^{(tr)}$*6hr*), and *resistance_status* indicates PC½ greater/smaller than 5hr. To discover the distinct transcriptional response for resistant and susceptible parasites individually, we performed comparative analysis for each parasite group (resistant/susceptible). We aimed to define top drug-response genes for each parasite group independent of age and batch effects (due to the collection and transportation issue with the $^{(bl)}$*0hr* and $^{(tr)}$*6hr* sample set, the treatment condition here unavoidably confounded with the batch of transcriptome measurement). To achieve that, we first extracted the expression residuals from the model (3) with the loess fit only which maximumly removed the expression variations caused by age from the raw data. Second, we conducted Mann-

Whitney test on the expression residuals between treatment conditions to compare 216 $^{(bl)}$*0hr* to 180 $^{(tr)}$*6hr* samples for resistant parasites and 168 $^{(bl)}$*0hr* to 130 $^{(tr)}$*6hr* samples for susceptible parasites with microarray measurement. To balance the sample size differences, subsampling was applied to 130 samples 100 times per gene to obtain the *p-value* at 80% confident level. For the multiple test correction, we calculated FDR for each gene using the distribution of 500, 000 null p-values generated from expression permutation based on all $^{bl)}$*0hr* and $^{(tr)}$*6hr* samples. To control the effect of treatment/batch, the structure of treatment condition was maintained during each time permutation.

We repeated the same analysis with the RNA-seq data. Mann-Whitney test was conducted to compare 67 $^{(bl)}$*0hr* to 55 $^{(tr)}$*6hr* samples for resistant parasites and 57 $^{(bl)}$*0hr* to 46 $^{(tr)}$*6hr* samples for susceptible parasites. And the *p-values* were obtained by 100 subsampling of 46 samples per gene. The result was merged with that from microarray for susceptible and resistance group, respectively, at FDR<0.05.

By this approach, we identified 20 significantly induced genes and 73 repressed genes upon drug in the KEL1PLA1 resistant parasites and 33 induced genes and 106 repressed genes in the WT susceptible parasites (FDR<0.05, corresponding *p*<1e-14). Significant common response genes were observed between the resistant and susceptible parasite samples which included up to 12 induced and 34 repressed genes (**Fig. 3b and Data S2**).

### References


1. R. W. van der Pluijm *et al.*, Triple artemisinin-based combination therapies versus artemisinin-based combination therapies for uncomplicated Plasmodium falciparum malaria: a multicentre, open-label, randomised clinical trial. *Lancet*, (2020).
2. M. Kucharski *et al.*, A comprehensive RNA handling and transcriptomics guide for high-throughput processing of Plasmodium blood-stage samples. *Malar. J.* **19**, 363 (2020).
3. S. Picelli *et al.*, Full-length RNA-seq from single cells using Smart-seq2. *Nat. Protoc.* **9**, 171-181 (2014).
4. Z. Bozdech *et al.*, The transcriptome of Plasmodium vivax reveals divergence and diversity of transcriptional regulation in malaria parasites. *Proc. Natl. Acad. Sci. U. S. A.* **105**, 16290-16295 (2008).
5. M. J. Mackinnon *et al.*, Comparative transcriptional and genomic analysis of Plasmodium falciparum field isolates. *PLoS Pathog.* **5**, e1000644 (2009).
6. M. E. Ritchie *et al.*, Limma powers differential expression analyses for RNA-sequencing and microarray studies. *Nucleic Acids Res.* **43**, e47 (2015).
7. D. Kim, B. Langmead, S. L. Salzberg, HISAT: a fast spliced aligner with low memory requirements. *Nat Methods* **12**, 357-360 (2015).
8. A. R. Quinlan, I. M. Hall, BEDTools: a flexible suite of utilities for comparing genomic features. *Bioinformatics* **26**, 841-842 (2010).
9. L. Zhu *et al.*, The origins of malaria artemisinin resistance defined by a genetic and transcriptomic background. *Nat Commun* **9**, 5158 (2018).
10. J. E. Lemieux *et al.*, Statistical estimation of cell-cycle progression and lineage commitment in *Plasmodium falciparum* reveals a homogeneous pattern of transcription in ex vivo culture. *Proc. Natl. Acad. Sci. U. S. A.* **106**, 7559-7564 (2009).
11. C. R. John *et al.*, M3C: Monte Carlo reference-based consensus clustering. *Sci. Rep.* **10**, 1816 (2020).

**Supplementary Figure Legend:**

**Table S1:** Summary of samples in this study.

Table S1

| | | $^{(bl)}$ *0hr* | | | $^{(tr)}$ *6hr* | | |
|---|---|---|---|---|---|---|---|
| | | Total | Microarray | RNA-seq | Total | Microarray | RNA-seq |
| | | 584 | 577 | 188 | 461 | 459 | 159 |
| Bangladesh | 1.Ramu | 115 | 110 | 47 | 91 | 91 | 43 |
| Myanmar | 2.Thabeikkyin | 22 | 21 | 7 | 15 | 15 | 7 |
| | 3.Pyin Oo Lwin | 12 | 12 | 3 | 11 | 11 | 2 |
| | 4.Pyay | 27 | 27 | 12 | 24 | 24 | 9 |
| | 5.Ann | 26 | 26 | 2 | 19 | 19 | 2 |
| Laos | 6.Sekong | 10 | 10 | 1 | 0 | 0 | 0 |
| Thai-Cambodia border | 7.Pusing | 31 | 31 | 12 | 30 | 30 | 13 |
| | 8.Khun Han | 6 | 6 | 0 | 3 | 3 | 0 |
| Cambodia | 9.Preah Vihear | 7 | 7 | 2 | 5 | 5 | 2 |
| | 10.Ratanakiri | 80 | 80 | 20 | 57 | 57 | 12 |
| | 11.Pailin | 57 | 57 | 32 | 51 | 51 | 30 |
| | 12.Pursat | 78 | 78 | 20 | 64 | 63 | 14 |
| Southern Vietnam | 13.Binh Phuoc | 113 | 112 | 30 | 91 | 90 | 25 |

### Supplementary Figures

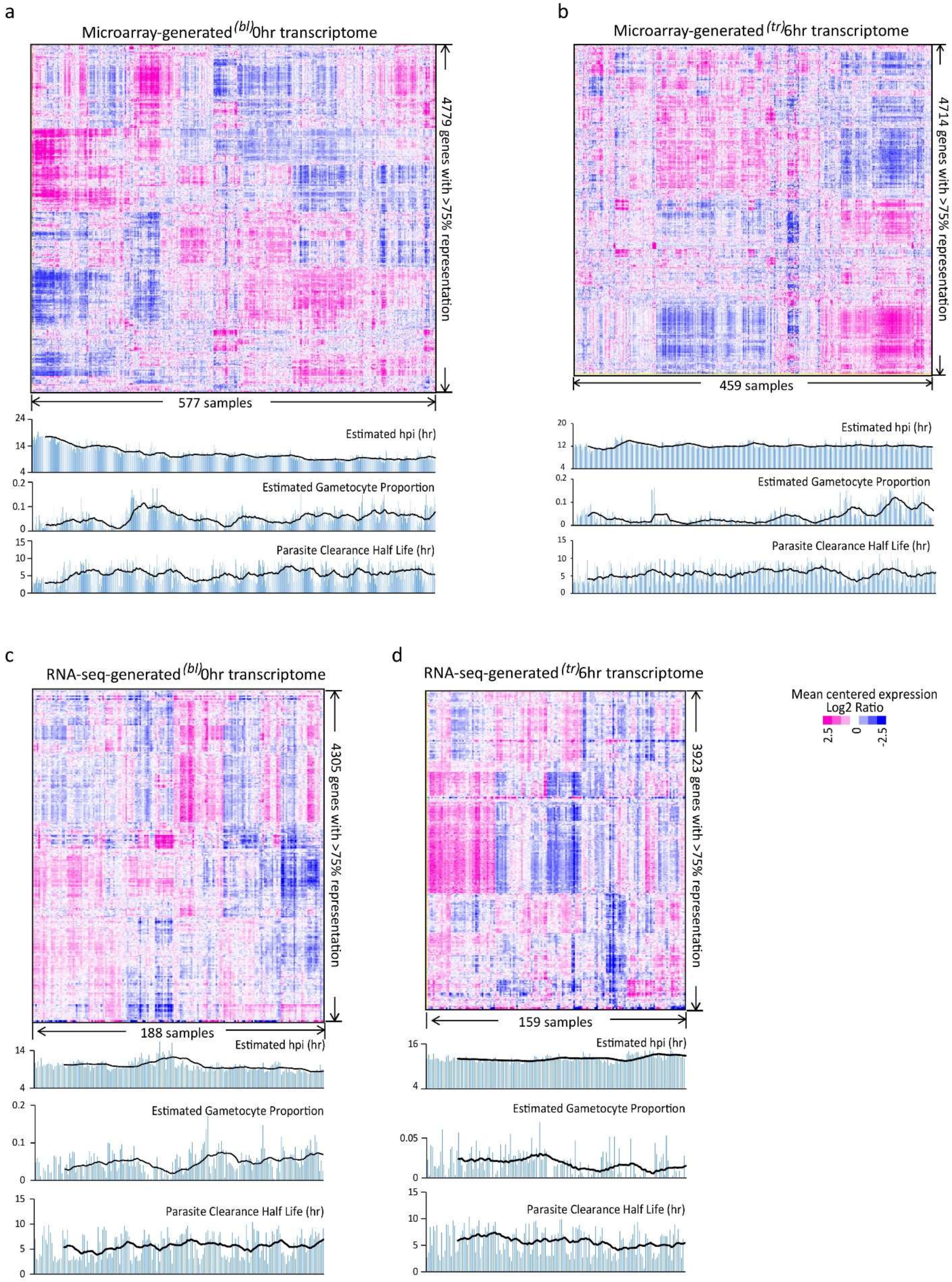

**Figure S1**: Transcriptional profiles of the studied *P.falciparum* parasites. Expression was normalized across samples by mean centering for each data set. Hierarchical clustering method was applied to the similarity matrix of samples/genes based on Pearson correlation coefficient. Only genes presented in >75% samples of a data set were used for the clustering analysis. a. Heat map of 577 $^{(bl)}0hr$ transcriptomes with 4779 representative genes obtained by microarrays. b. Heat map of 459 $^{(tr)}6hr$ transcriptomes with 4714 representative genes obtained by microarrays. c. Heat map of 188 $^{(bl)}0hr$ transcriptomes with 4305 representative genes obtained by RNA-seq. d. Heat map of 159 $^{(tr)}6hr$ transcriptomes with 3923 representative genes obtained by RNA-seq.

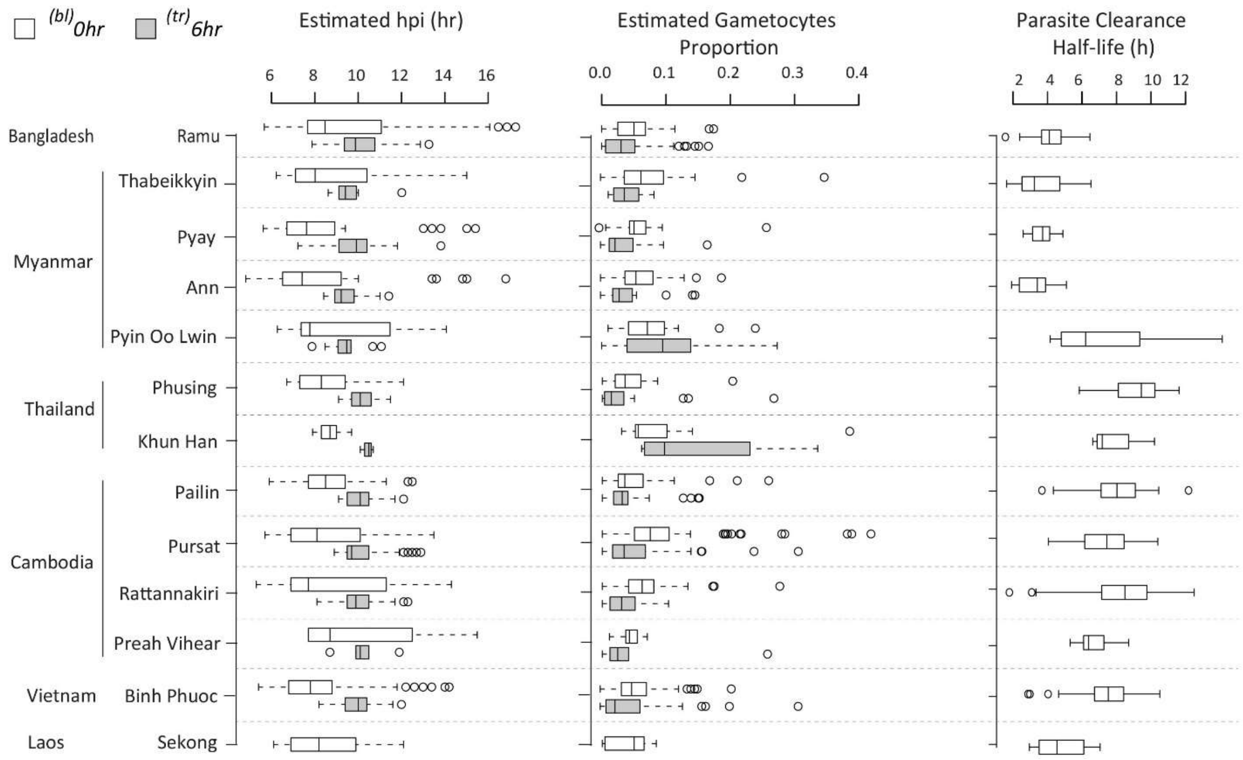


**Figure S2**: Distribution of the estimated age/hpi, gametocytes fraction and PC½ across the sampling sites listed by country. The data was shown for all the samples before transcriptome filtering. White boxes represent $^{(bl)}$*0hr* samples and grey boxes represent $^{(tr)}$*6hr* samples.

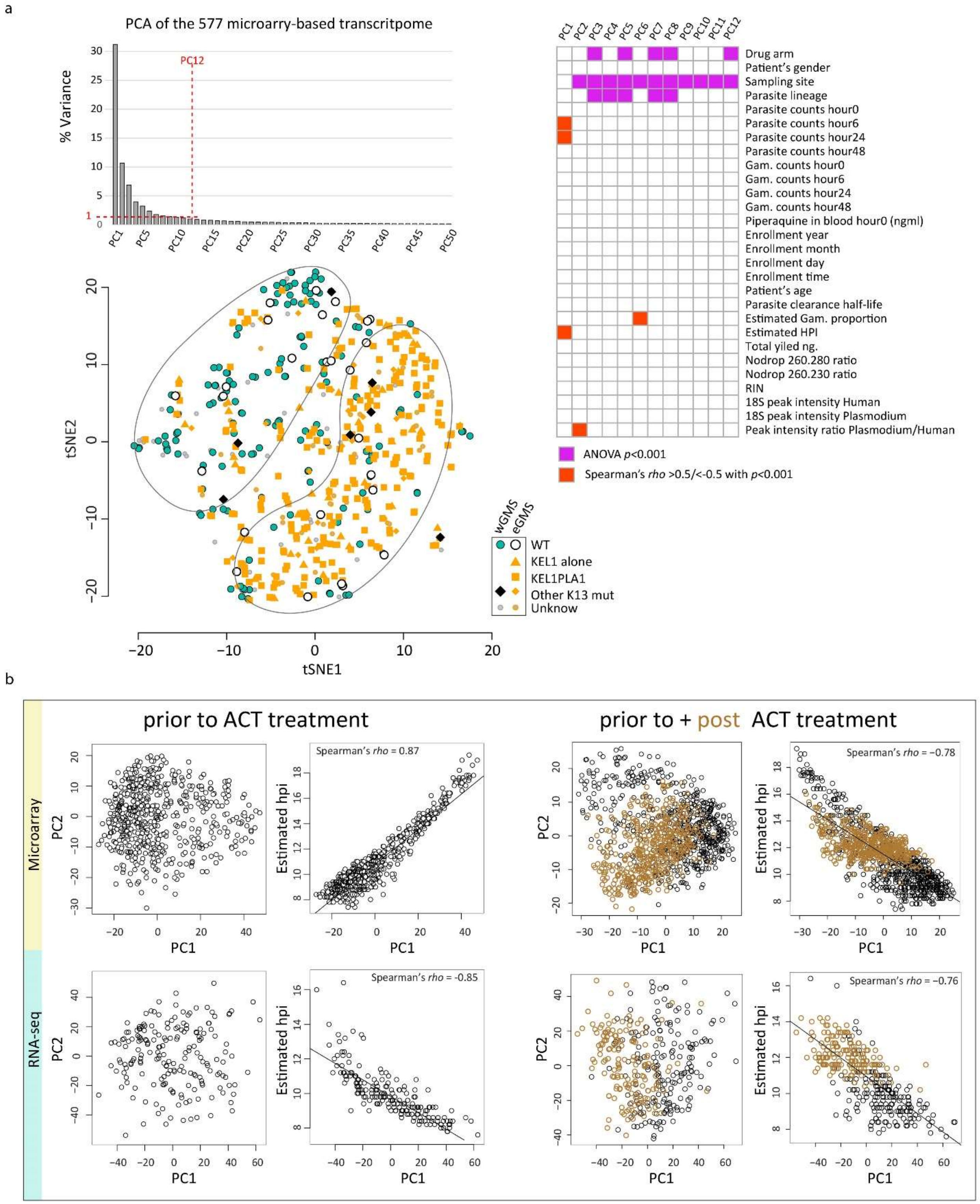
a
PCA of the 577 microarry-based transcritpome
% Variance
PC12
tSNE1
tSNE2
wGMS
eGMS
WT
KEL1 alone
KEL1PLA1
Other K13 mut
Unknow
Drug arm
Patient's gender
Sampling site
Parasite lineage
Parasite counts hour0
Parasite counts hour6
Parasite counts hour24
Parasite counts hour48
Gam. counts hour0
Gam. counts hour6
Gam. counts hour24
Gam. counts hour48
Piperaquine in blood hour0 (ngml)
Enrollment year
Enrollment month
Enrollment day
Enrollment time
Patient's age
Parasite clearance half-life
Estimated Gam. proportion
Estimated HPI
Total yiled ng.
Nodrop 260.280 ratio
Nodrop 260.230 ratio
RIN
18S peak intensity Human
18S peak intensity Plasmodium
Peak intensity ratio Plasmodium/Human
ANOVA p<0.001
Spearman's rho >0.5/<-0.5 with p<0.001
b
prior to ACT treatment
prior to + post ACT treatment
Microarray
RNA-seq
Spearman's rho = 0.87
Spearman's rho = −0.78
Spearman's rho = -0.85
Spearman's rho = −0.76
Estimated hpi
PC1
PC2

**Figure S3**: PCA of the parasite transcriptome. a. Bar plot represents the percent of variance explained by each PC. For the top 12 PCs, each explained >1% of the total transcriptome variance. The relation to clinical or technical factors were tested for each of the 12 PCs. The grid mixing PCs and factors represents the significant association in magenta for the PC-factor pairs passing the threshold of $p$<0.001 with ANOVA test and in red for the PC-factor pairs passing the threshold $p$<0.001 and Spearman's *rho*>0.5. The scatter plot represents the tSNE derived two-dimensional visualization of 577 $^{(bl)}$0hr samples by lineages and geographical regions. b. PC2 and estimated age/hpi were plotted against PC1 individually for each data set ($^{(bl)}$0hr or $^{(tr)}$6hr) by each technology (microarray or RNA-seq).

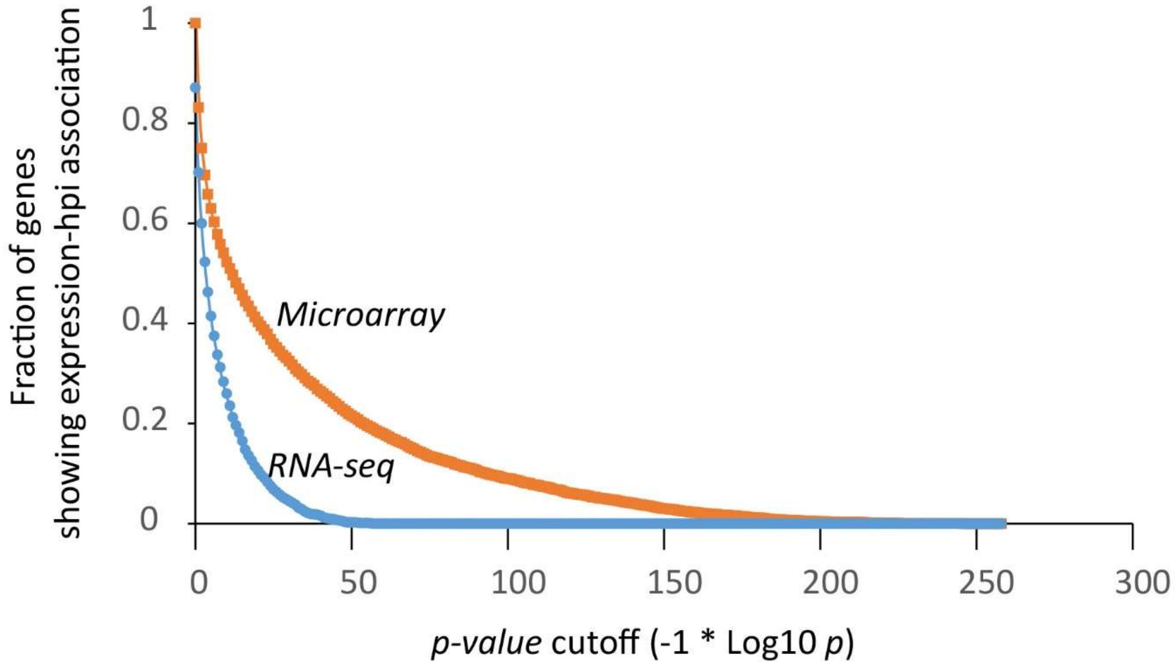


**Figure S4:** Expression level correlates to age/hpi for most of the *P. falciparum* genes. P-values were obtained by testing the null hypothesis of no expression change over hpi for each gene using the package gam of R. The curves plotted for Microarray data (orange) and RNA-seq data(blue) to show the gene fractions at each p-value cutoff for defining hpi-associated gene expression.

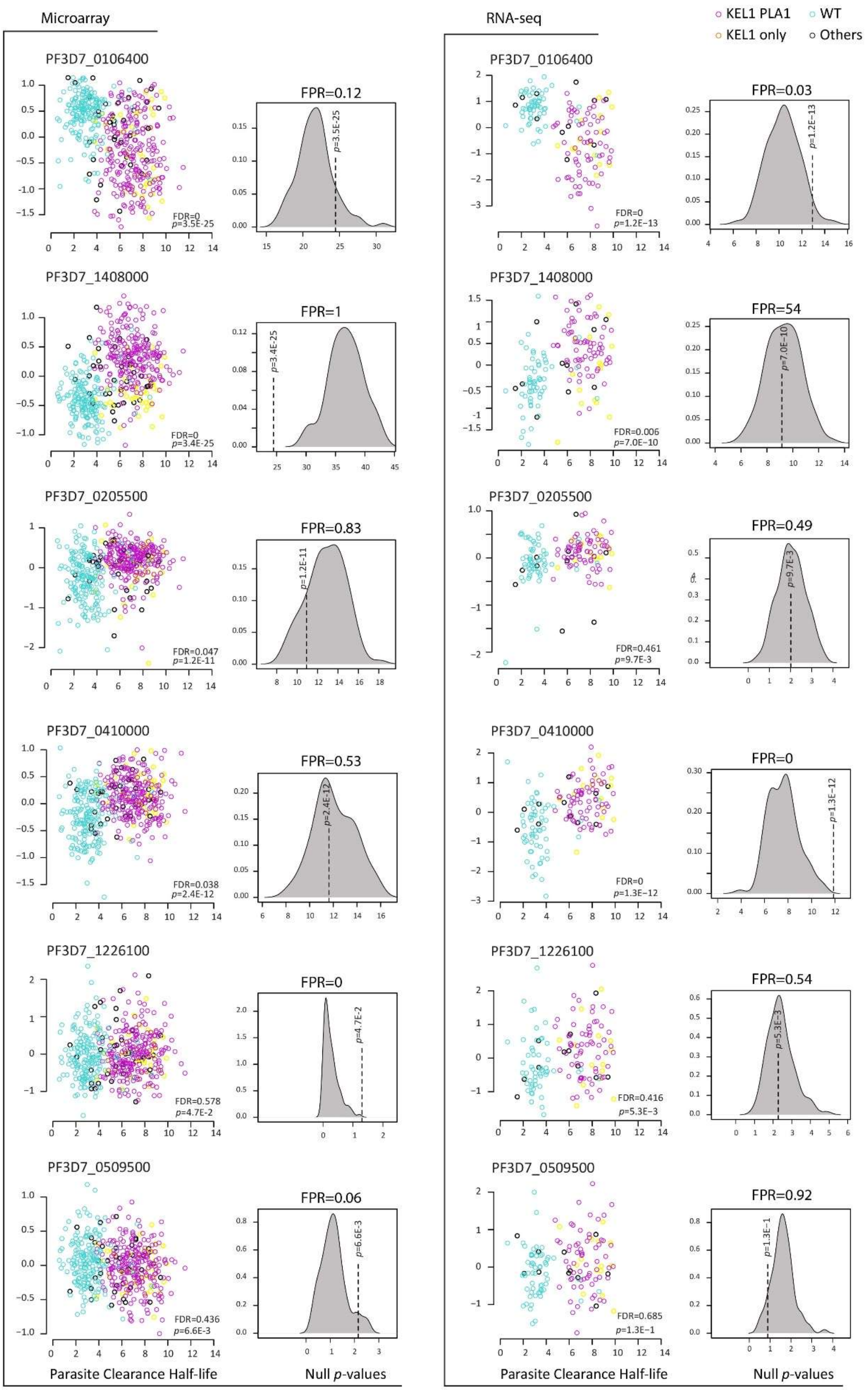

Microarray
RNA-seq
KEL1 PLA1
WT
KEL1 only
Others
PF3D7_0106400
FPR=0.12
p=3.5E-25
FDR=0
p=3.5E-25
PF3D7_0106400
FPR=0.03
p=1.2E-13
FDR=0
p=1.2E-13
PF3D7_1408000
FPR=1
p=3.4E-25
FDR=0
p=3.4E-25
PF3D7_1408000
FPR=54
p=7.0E-10
FDR=0.006
p=7.0E-10
PF3D7_0205500
FPR=0.83
p=1.2E-11
FDR=0.047
p=1.2E-11
PF3D7_0205500
FPR=0.49
p=9.7E-3
FDR=0.461
p=9.7E-3
PF3D7_0410000
FPR=0.53
p=2.4E-12
FDR=0.038
p=2.4E-12
PF3D7_0410000
FPR=0
p=1.3E-12
FDR=0
p=1.3E-12
PF3D7_1226100
FPR=0
p=4.7E-2
FDR=0.578
p=4.7E-2
PF3D7_1226100
FPR=0.54
p=5.3E-3
FDR=0.416
p=5.3E-3
PF3D7_0509500
FPR=0.06
p=6.6E-3
FDR=0.436
p=6.6E-3
PF3D7_0509500
FPR=0.92
p=1.3E-1
FDR=0.685
p=1.3E-1
Parasite Clearance Half-life
Null p-values
Parasite Clearance Half-life
Null p-values

**Figure S5:** The expression residuals were plotted against the PC½ values with parasite lineage of KEL1PLA1 indicated by purple circles, KEL1 only by yellow, WT by turquoise and others by black for selected genes: TSR2 (PF3D7_0106400), PMII(PF3D7_1408000), PF3D7_0205500 and EVP1(PF3D7_0410000), HAD3(PF3D7_1226100) and PF3D7_1467000 which showed different levels of FDR and FPR. The density plot on the right represents the null *p*-values distribution for FPR calculation based on 100 times permutation of each corresponding gene's resistance status/PC½ values within lineages.

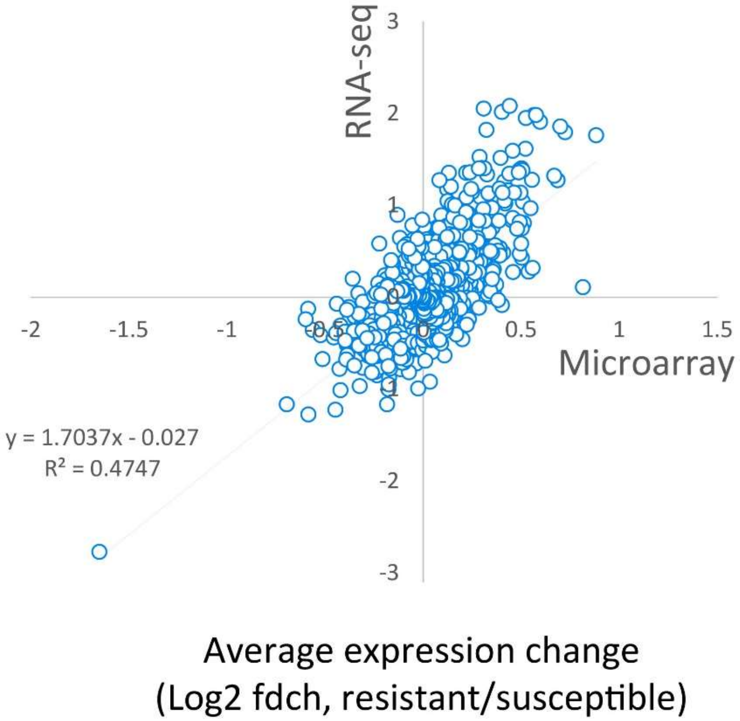


**Figure S6:** The average expression change determined by RNA-seq was plotted against that by microarray to show the consistent results of TWAS analysis regardless of the methods applied. The average expression change was calculated as the difference between the average transcriptional level (Log2 Ratio) of a gene in the resistant parasites (PC½>5hr) and that in the susceptible parasites (PC½<5hr). Therefore, the average expression change represented here are Log2 fold change of transcriptional level (Log2 fdch).

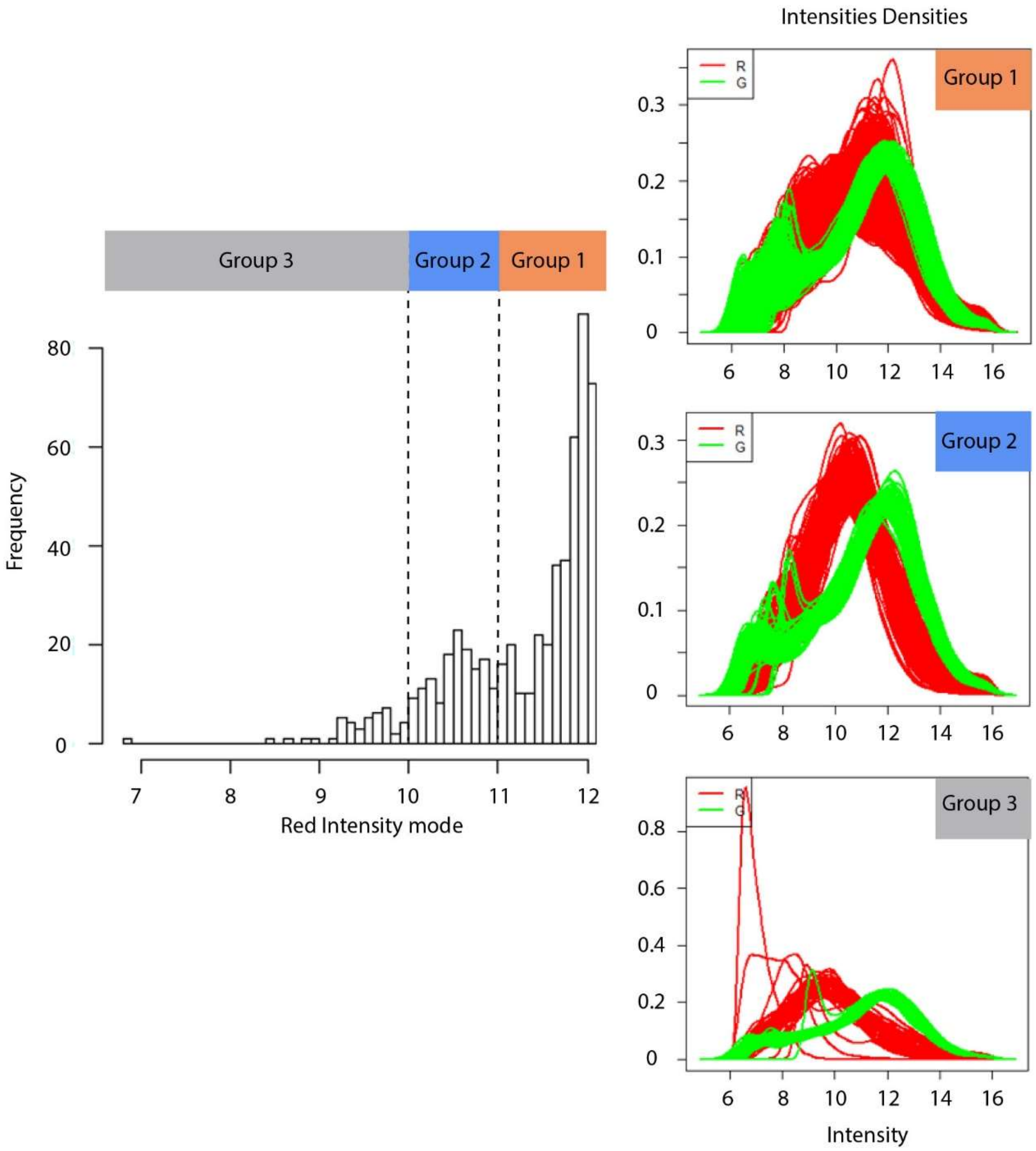


**Figure S7**: Defined intensity threshold for microarray-generated transcriptome filtering using the *(tr)6hr* data set. The histogram on the left represents the overall distribution of red intensity mode values for all the 659 studied arrays. The mode value is the most frequently appearing value in a data set. Here, we estimated it using the intensity value appearing at the biggest peak of corresponding density plot. Arbitrary cutoffs were set at the mode value of 10 and 11 to bin the samples/arrays into 3 groups. The density plot of red intensity (Cy5) and green (Cy3) intensity is drawn for each group on the right. The threshold was set at the mode value of 10 to select samples displaying sufficient signals for the subsequent analysis.
